# Adrenal cortex size, homeostasis and tumorigenesis is regulated by gonadal hormones via androgen receptor/β-catenin signalling crosstalk

**DOI:** 10.1101/2022.06.23.497219

**Authors:** Rodanthi Lyraki, Anaëlle Grabek, Amélie Tison, Mirko Peitzsch, Nicole Bechman, Sameh A Youssef, Alain de Bruin, Elvira R.M. Bakker, Frank Claessens, Marie-Christine Chaboissier, Andreas Schedl

## Abstract

Female bias is highly prevalent among adrenal cortex hyperplasia and neoplasia, but the reasons behind this phenomenon are poorly understood. In this article, we show that overexpression of the secreted WNT agonist R-spondin-1 leads to ectopic activation of WNT/β-catenin signalling and causes sex-specific adrenocortical hyperplasia in mice. While female adrenals show ectopic proliferation, male adrenals display excessive immune system activation and cortical thinning. Using a combination of genetic manipulations and hormonal treatment, we show that gonadal androgens suppress ectopic proliferation in the adrenal cortex and determine the selective activation of WNT-related genes *Axin2* and *Wnt4*. Notably, genetic removal of androgen receptor (AR) from adrenocortical cells restores the mitogenic effect of WNT/β-catenin signalling. This is the first demonstration that AR activity in the adrenal cortex determines susceptibility to canonical WNT signalling-induced hyperplasia.

**Teaser:** Activation of R-spondin signaling in the adrenal cortex leads to a sexually dimorphic phenotype causing tumors in females and immune cell recruitment in males

## Introduction

Sexual dimorphism is prevalent among mammalian phenotypic traits ^1^ and underlies several aspects of mammalian physiology including malignant transformation ^2^ and immunity ^3^. Sex-specific effects often stem from the action of gonadal hormones ^4^ but can also have sex chromosome-related causes, such as the incomplete inactivation of X chromosome genes ^5^. An important open question is whether sex impacts size maintenance and homeostasis of self-renewing adult tissues, such as the adrenal cortex.

Adrenals are endocrine organ consisting of the inner medulla, responsible for the synthesis of catecholamines, and the outer cortex, responsible for the synthesis of steroid hormones. The adrenal cortex is characterised by the expression of the transcription factor SF1 (*Nr5a1*) and is further divided into concentric rings that form a characteristic zonation pattern ^6^. The outer zona glomerulosa (zG) produces mineralocorticoids, the middle zona fasciculata (zF) produces glucocorticoids, and the inner zona reticularis (zR) produces androgens. The latter is absent in mice; in its place, we find the X-zone, a transient remnant of the fetal adrenal with unknown functions in adulthood ^7^. Finally, the whole organ is surrounded by the non-endocrine capsule, composed of a mesothelial layer enclosing cells of mesenchymal origin.

The adrenal cortex undergoes constant renewal thanks to resident populations of stem/progenitor cells that are primarily concentrated in the capsule and the sub-capsular zG ^8–11^. A proliferating zone can be distinguished in the outer cortex that gradually fades out towards the inner part; as a result, the inner zF is largely composed of quiescent cells ^12^. Proliferation arrest coincides with transdifferentiation of zG cells to a zF identity and centripetal migration ^13^. Finally, older cells are eliminated at the corticomedullary border, presumably via apoptosis.

Canonical WNT/β-catenin signalling has a prominent position among the molecular pathways that participate in maintaining adrenal cortex homeostasis and zonation ^14,15^. Dramatic interventions such as the constitutive activation of β-catenin lead to the expansion of the zG to the expense of the zF and tumour development in ageing mice (Berthon et al. 2010;, Pignatti et al. 2020). Other mechanisms allow for a more precise fine-tuning of WNT activation levels in the adrenal cortex. A negative feedback loop attenuates WNT signalling based on the activity of the membrane-bound E3 ubiquitin ligase ZNRF3, which mediates the ubiquitin-dependent endocytosis and lysosomal degradation of Frizzled (FZD) WNT receptors ^18^. Secreted ligands of the R-spondin family (RSPO) and their cognate receptors LGR4/5/6 form a complex that can bind and remove ZNRF3 from the cell surface, thus potentiating WNT signalling ^19,20^. In the mouse adrenal cortex, a diminishing gradient of WNT signalling activity from the zG to the zF is maintained via the localised expression of secreted WNT potentiators, mainly *Wnt4* in the zG, and R-spondins (*Rspo1* and *Rspo3*) in the capsule ^21–23^. The R-spondin/LGR/ZNRF3 axis creates a balance that ensures the maintenance of organ size. While *Rspo3* deletion in mice results in cortical atrophy ^21^, *Znrf3* deletion leads to adrenal hyperplasia ^22^. Although this knowledge originates mostly from mouse genetic studies, the high prevalence of *CTNNB1* and *ZNRF3* driver mutations in human adrenocortical carcinoma (ACC) shows the relevance of these pathways for human adrenal disease ^24,25^.

The adrenal gland is recognised as one of the most sexually dimorphic non-reproductive organs. For example, many forms of adrenocortical hyperplasia and neoplasia associated with endocrine manifestations, such as Cushing’s syndrome, are more frequent among women than men ^26^. This includes benign adrenocortical adenomas (female:male ratio 4-8:1) ^27,28^, and ACC (female:male ratio 1.5-2.5:1) ^29–31^. Under normal homeostatic conditions, the mouse adrenal shows a strong dimorphism, with female adrenals being significantly bigger than males and consistently displaying higher proliferation levels ^32,33^. Moreover, we recently showed that adrenocortical renewal is more rapid in female than in male mice, due to higher activity of cortical AXIN2^+^ progenitors and female-specific recruitment of capsular GLI1^+^ progenitors which is dramatically affected by gonadectomy and the administration of testicular androgens ^33^. Finally, it has been reported that androgens influence the adrenal cortex by enhancing WNT signalling and antagonising PKA activity ^34^.

Even though testicular androgens influence the adrenal cortex, whether this influence is direct, and how this influence translates to a reduced susceptibility to hyperplasia is still obscure. To answer these questions, we used a mouse model of disrupted adrenal homeostasis due to the ectopic expression of *Rspo1* in the adrenal cortex, thus causing moderate WNT signalling hyperactivation. This genetic manipulation results in ectopic proliferation and hyperplasia in female mice, in contrast to cortical thinning and degeneration in males. We show that sexual dimorphism in our model is dependent on testicular androgens, which act directly on adrenocortical cells through their cognate receptor AR to cause cell cycle arrest and counteract the mitogenic effect of enhanced WNT signalling.

## Results

### Ectopic expression of *Rspo1* leads to sex-specific adrenocortical hyperplasia or degeneration

In order to generate a mouse model for adrenocortical hyperplasia, we sought to disrupt the gradient of canonical WNT signalling activation by ectopically expressing R-spondin 1 (RSPO1) in the adrenal cortex. We used a Cre-inducible *Rspo1* gain-of-function (GOF) allele ^35,36^ and the *Sf1-cre* transgene that drives Cre recombinase expression in SF1(+) tissues including the adrenal cortex ^37^ (*Sf1-Rspo1^GOF^* mice) (Figure 1A). In control adult animals *Rspo1* expression was restricted to the outer adrenal capsule, in agreement with previous research ^21^. By contrast, *Sf1-Rspo1^GOF^* mice showed expression throughout the adrenal cortex (Figure 1B). Consistent with the dramatically expanded expression domain, *Rspo1* mRNA levels were on average 70-80-fold higher in *Sf1-Rspo1^GOF^* adrenals compared to controls. No significant differences were detected between males and females (Figure S1A).

**Figure 1.**
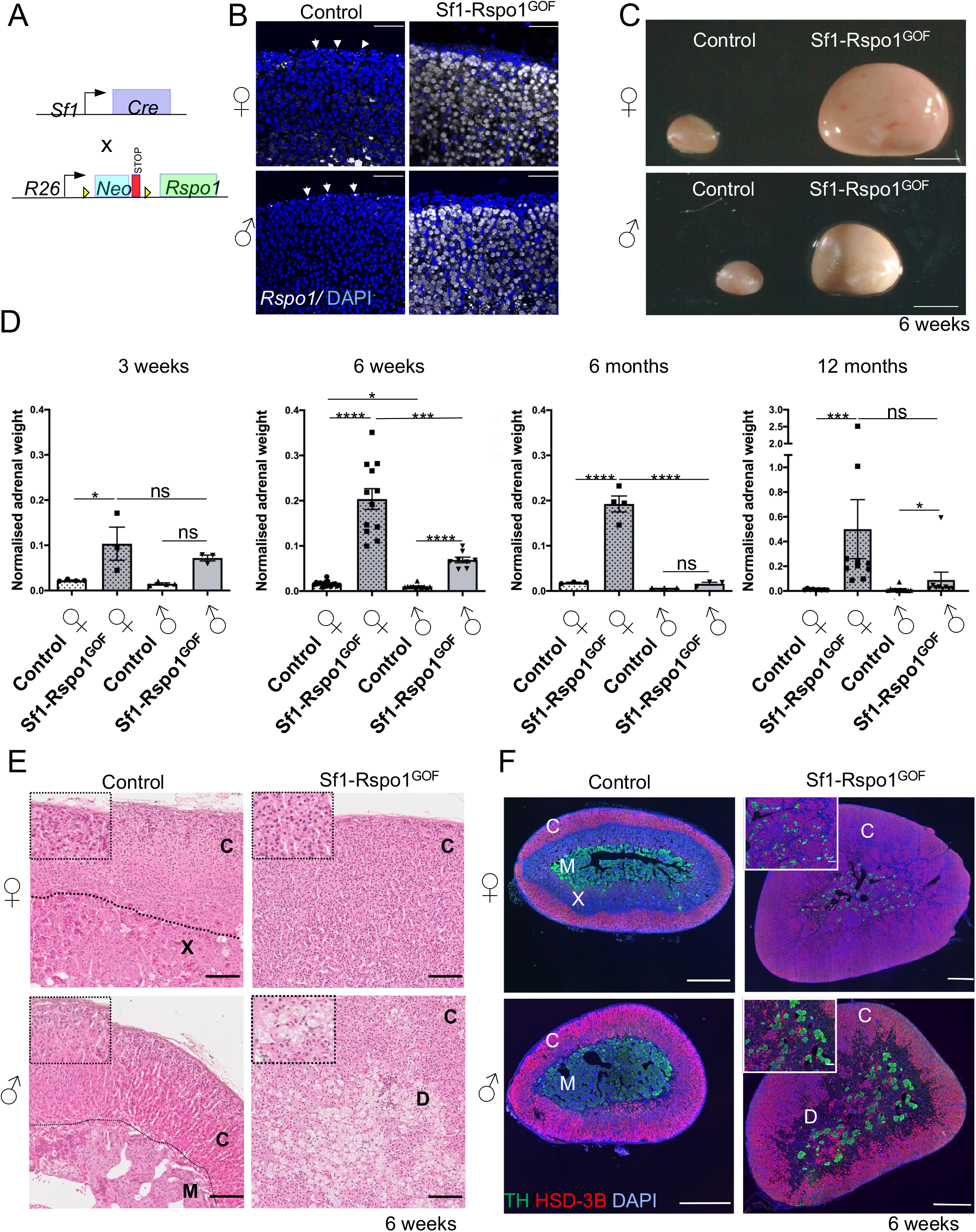
RSPO1 overexpression leads to sex-specific adrenocortical hyperplasia or degeneration. **A**) Schematic representation of the genetic strategy to overexpress RSPO1 in steroidogenic cells using the Sf1-Cre. **B**) In situ hybridisation for *Rspo1* using RNA Scope technology on adrenal sections from 3 month-old mice. White arrows mark *Rspo1* mRNA molecules. Scale bars: 50 μm. **C**) Representative photographs of adrenal glands from control and *Sf1-Rspo1^GOF^* mice at the end of puberty (6 weeks of age). Scale bars: 1mm. **D**) Graphs of mean adrenal weight normalised to body weight at different ages comparing control mice to *Sf1-Rspo1^GOF^* mice when ectopic RSPO1 expression driven by Sf1-Cre (error bars represent SEM). Statistical analysis for 3 weeks and 6 months was performed using ordinary one-way ANOVA followed by Tukey’s multiple comparisons test. Statistical analysis for 6 weeks was performed using Welch’s one-way ANOVA followed by Dunnett’s T3 post-hoc test. Statistical analysis for 12 months was done using Kruskal Wallis test. Adjusted P values for 3-weeks comparisons: Control F vs *Sf1-Rspo1^GOF^* F: P=0.0192, Control M vs *Sf1-Rspo1^GOF^* M: P=0.1067, *Sf1-Rspo1^GOF^* F vs *Sf1-Rspo1^GOF^* M: P=0.5631. Adjusted P values for 6 weeks comparisons: Control F vs *Sf1-Rspo1^GOF^* F: P<0.0001, Control F vs control M: P=0.0239, *Sf1-Rspo1^GOF^* F vs *Sf1-Rspo1^GOF^* M: P=0.0005, Control M vs *Sf1-Rspo1^GOF^* M: P<0.0001. Adjusted P values for 6 months comparisons: Control F vs *Sf1-Rspo1^GOF^* F: P<0.0001, Control M vs *Sf1-Rspo1^GOF^* M: P=0.9500, *Sf1-Rspo1^GOF^* F vs *Sf1-Rspo1^GOF^* M: P=0.0082. Adjusted p-values for 12 months comparisons: Control F vs *Sf1-Rspo1^GOF^* F: P=0.0008, Control M vs *Sf1-Rspo1^GOF^* M: P=0.0144, *Sf1-Rspo1^GOF^* F vs *Sf1-Rspo1^GOF^* M: P=0.4853. **E**) H&E staining of adrenals from 6 week-old mice. C: cortex, X: x-zone, M: medulla, D: degeneration. Scale bar: 100 μm, inserts: 20 μm. **F**) Immunofluorescence staining for Tyrosine hydroxylase (TH, marker of the adrenal medulla) and 3βHSD (marker of steroidogenic cells), using adrenal sections from 6 week-old mice. C: cortex, X: x-zone, M: medulla, D: degeneration. Note that the size of the scale bar is different for each picture. Scale bar: 500 μm, inserts: 100 μm. M: Male, F: female.

Ectopic *Rspo1* expression in our models resulted in striking hyperplasia of the adrenal glands in 6 weeks old mice (Figure 1C). Because the *Sf1-Cre* line drives expression of the GOF allele already during embryogenesis, hyperplasia was noticeable in pre-pubertal pups that was comparable between male and female mice at 3 weeks of age (Figure 1D). After puberty, however, the phenotype evolved in a highly sexually dimorphic manner. Whereas female adrenals from *Sf1-Rspo1^GOF^* mice further increased in size, male adrenals remained smaller, and their size even regressed as they aged (Figure 1D, 6 weeks, 6 months, and 12 months). Thus, puberty appears to be a critical period for the development of sexual dimorphism in our model.

Next, we conducted a histological analysis to examine the cellular composition of *Sf1-Rspo1^GOF^* adrenals. According to H&E analysis at 6 weeks of age, all the female GOF adrenals analysed exhibited diffuse atypical hyperplasia in the cortex (n=5) (Figure 1E). The inner cortex was composed of steroidogenic cells expressing 3βHSD, although its expression seemed reduced compared to control adrenals, while the medulla was fragmented (Figure 1F). Male GOF adrenals, on the other hand, displayed non-neoplastic degenerative changes in the form of markedly vacuolated, polynucleated cells. At 6 weeks, all the male *Sf1-Rspo1^GOF^* adrenals exhibited these degenerative changes to a varying degree, ranging from a few abnormal cells to extensive degenerated lesions (n=6 mice analysed) (Figure 1E) which were negative for the steroidogenic marker 3βHSD (Figure 1F). By 3 months of age, degenerative areas expanded significantly in all the male *Sf1-Rspo1^GOF^* adrenals (n=3), leading to cortical thinning (Figure S1B). These degenerative changes affected female adrenals to a much lesser degree (2/3 adrenals examined at 3 months displayed only a few abnormal cells) (Figure S1B). zG expansion is a known consequence of the constitutive activation of β-catenin in the adrenal cortex ^16,17^. However, in *Sf1-Rspo1^GOF^* adrenals, the zG was not expanded in either sex, as shown by immunostaining with the zG markers DAB2 and LEF1 (Figure S1C,D).

To assess whether ectopic expression of R-spondin affected the endocrine activity of the hyperplastic adrenals, we measured plasma steroids of control and GOF animals by LC-MS/MS. The mineralocorticoid hormone aldosterone and its precursor 18-OH-corticosterone were not significantly changed among our experimental groups. Levels of the glucocorticoid corticosterone and another steroid hormone precursor, 11-deoxycorticosterone, showed a mild decrease in female and increase in male *Sf1-Rspo1^GOF^* mice, although none of these changes reached statistical significance (Figure S1F). Analysis of the adrenocorticotropic hormone (ACTH)-responsive differentiation marker AKR1B7 ^38^ showed loss of expression in subpopulations of cells of the inner cortex, which explains the absence of hormone overproduction despite the striking hyperplasia (Figure S1E).

As the *Sf1-Rspo1^GOF^* animals age, the diffuse hyperplasia and cortical thinning give way to increased frequency of well-circumcised benign nodules and adenomas (Figure S2) (6/8 *Sf1-Rspo1^GOF^* males and 6/9 *Sf1-Rspo1^GOF^* females of 12 months). Importantly, 1 out of 8 aging *Sf1-Rspo1^GOF^* females had a well differentiated adrenal carcinoma with capsular invasion (Figure S2). None of the controls or male *Sf1-Rspo1^GOF^* animals displayed any malignant tumour suggesting sexual dimorphism also with regard to tumour progression.

Taken together, ectopic *Rspo1* expression exerts a highly sexually dimorphic effect on the adrenal cortex: In female mice, it leads to diffuse hyperplasia of the adrenal cortex without increased endocrine activity. On the contrary, male GOF adrenals display expansive degenerative lesions and cortical thinning, while compensating activity of the remaining cortex ensures that insufficiency of steroid hormones is avoided.

### Ectopic expression of *Rspo1* leads to female-specific ectopic proliferation

The striking sexual dimorphism in our *Sf1-Rspo1^GOF^* model is reminiscent of the female bias observed in human adrenal diseases. To further investigate its molecular underpinnings, we conducted mRNA sequencing and differential expression analysis of whole adrenals from control and *Sf1-Rspo1^GOF^* mice of both sexes during puberty (at 4 weeks), a time point before male specific degeneration of adrenals occurs (Figure S3A; data available at NCBI’s Gene expression Omnibus ^39^ with GEO Series accession number GSE178958). As expected, principal component analysis (PCA) identified the presence of the *Rspo1^GOF^* allele as a major component in our RNA-Seq experiment. Strikingly, sex was the second most important factor underlying the variation in gene expression patterns among our experimental groups (Figure 2A). To gain insights into the molecular changes occurring in different subgroups we next performed gene set enrichment analysis using the Molecular Signature Database (Broad Institute). Genes enriched in *Sf1-Rspo1^GOF^* animals of both sexes compared to controls (corresponding to cluster 1 of the heatmap in Figure S3B) were related to β-catenin upregulation ^40,41^ or targets of the β-catenin -associated transcription factor LEF1 ^42^, a finding consistent with an hyperactivation of canonical WNT signalling by *Rspo1*. Genes specifically upregulated in female *Rspo1^GOF^* adrenals (corresponding to cluster 3 of the heatmap in Figure S3B) were found to be primarily related to cell cycle regulation, DNA replication and repair, and cell division (Figure 2B). A more in-depth gene set enrichment analysis revealed that targets of transcription factors belonging to the E2F family and the DREAM complex ^43^, critical repressors of cell cycle genes participating in the G1/S and G2/M transitions, were specifically upregulated in female *Sf1-Rspo1^GOF^* adrenals (Figure 2C). On the contrary, genes highly expressed in male *Sf1-Rspo1^GOF^* adrenals (corresponding to cluster 2 of the heatmap in Figure S3B) were associated with the regulation of immune system and defence response (Figure 2B). Moreover, an unbiased comparison of differentially regulated genes between male and female *Sf1-Rspo1^GOF^* adrenals via GSEA analysis confirmed that DNA replication and immune response were among the top enriched pathways in female and male *Sf1-Rspo1^GOF^* adrenals, respectively (Figure S3C). Of note, GSEA analysis revealed a downregulation of catecholamine secretion in *Sf1-Rspo1^GOF^* adrenals compared to controls, probably reflecting the fragmentation of the medulla that we observed in our histological analysis (Figure 1F).

**Figure 2.**
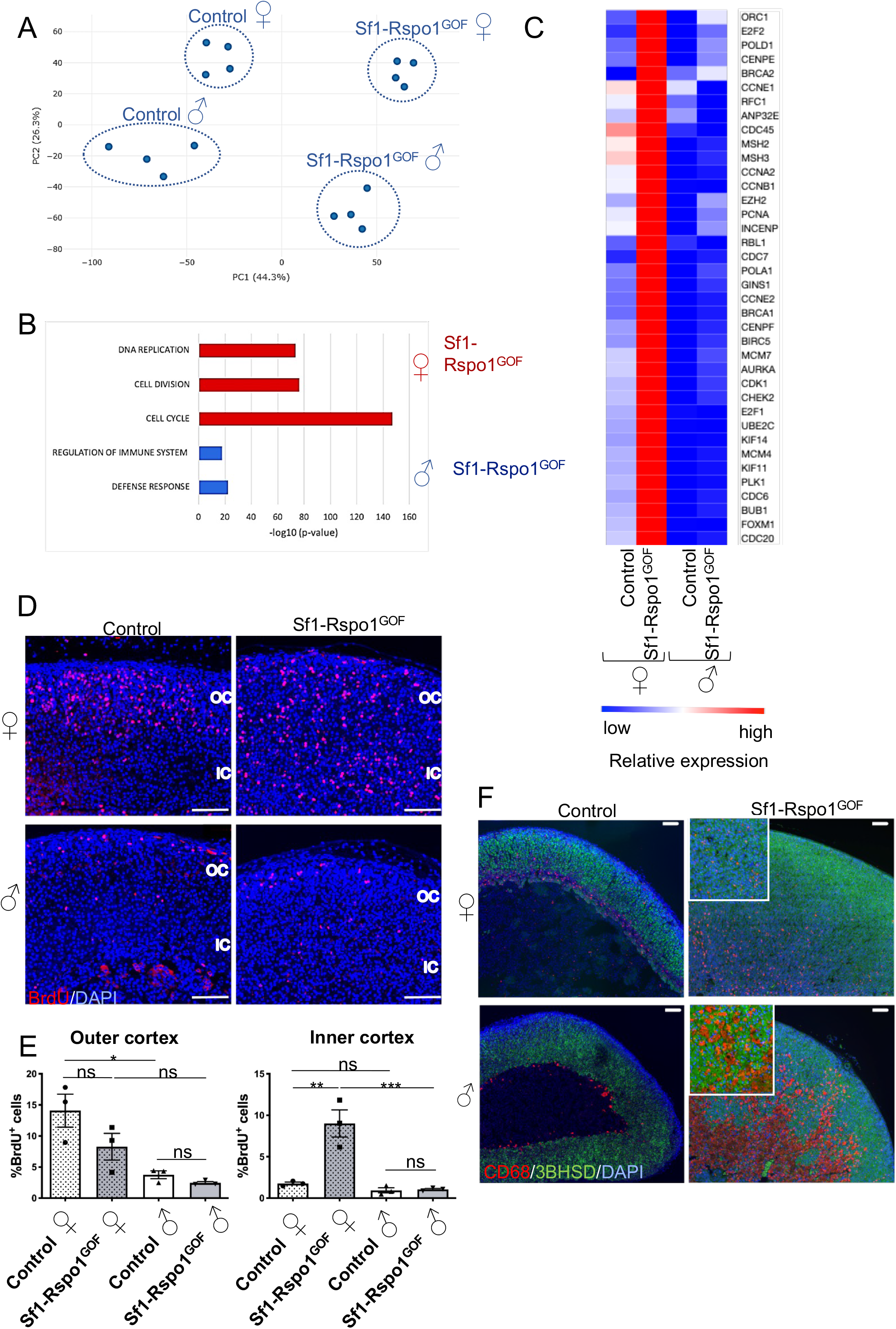
Transcriptomic analysis of *Sf1-Rspo1^GOF^* adrenals during puberty reveals sex-specific regulation of cell cycle and immune responses. **A**) Principal components analysis (PCA) plot of gene expression data in control and *Sf1-Rspo1^GOF^* male versus female adrenals during puberty. Every dot corresponds to an independent biological replicate. **B**) Heatmap representation of relative expression differences for proliferation genes among experimental groups. Relative expression among the groups is shown in colour code. The expression of all genes shown here is known to be induced during either the G1/S or the G2/M transitions of the cell cycle. **C**) Top enriched GO terms in the clusters representing genes that are highly expressed in male or female *Sf1-Rspo1^GOF^* adrenals compared to the other experimental groups. P-value indicates statistical significance. **D**) BrdU proliferation analysis shown as mean percentage of proliferating cells over total number of cells in the adrenal cortex of 6-week old mice (error bars represent SEM) (n=3). The area close to the capsule (outer cortex) is distinguished from the deeper layers (inner cortex). Statistical analysis was performed using ordinary one-way ANOVA followed by Tukey’s multiple comparisons test. Adjusted P-values for outer cortex graph: Control F vs *Sf1-Rspo1^GOF^* F: P=0.1623, Control F vs Control M: P=0.0128, *Sf1-Rspo1^GOF^* F vs *Sf1-Rspo1^GOF^* M: P=0.1638, Control M vs *Sf1-Rspo1^GOF^* M: P=0.9506. Adjusted P-values for inner cortex graph: Control F vs *Sf1-Rspo1^GOF^* F: P=0.0013, Control F vs Control M: P=0.8967, *Sf1-Rspo1^GOF^* F vs *Sf1-Rspo1^GOF^* M: P=0.0007, Control M vs *Sf1-Rspo1^GOF^* M: P=0.9996. **E)** Representative immunofluorescence images for BrdU. Scale bar: 100 μm. OC: outer cortex, IC: inner cortex. **F**) Immunofluorescence staining for CD68 (marker of murine macrophages) and 3βHSD (marker of steroidogenic cells), using adrenal sections from 6 week-old mice. White arrows point to macrophages within the adrenal cortex. Scale bar: 100 μm. M: male, F: female.

To confirm the conclusions drawn from the mRNA sequencing experiment, we analysed DNA replication in the adrenal cortex at 6 weeks of age via BrdU incorporation (Figure 2 D, E). No differences in proliferation between control and GOF adrenals were observed in the outer cortex, which makes up the proliferating zone under wild-type circumstances. However, proliferation was dramatically increased in the inner cortex of *Sf1-Rspo1^GOF^* females, which mostly consists of quiescent cells in the control animals. Surprisingly, no increase was observed in male *Sf1-Rspo1^GOF^* adrenals. Sex specific proliferation was not observed before puberty, as male and female GOF adrenals both show increased proliferation rates during embryogenesis (Figure S4 A,B).

Previous work from our group has shown the dominant role of RSPO3 over RSPO1 in the mouse adrenal cortex. Even though *Rspo3* deletion led to cortical atrophy both during development and in adulthood, *Rspo1* deletion did not produce an apparent defect ^21^. In order to test if the phenotype we observe is specific to *Rspo1^GOF^*, we analysed the effect of ectopic *Rspo3* expression in the adrenal cortex, taking advantage of a previously published *Rspo3^GOF^* allele in the *Rosa26* locus ^44^ and the *Sf1-Cre* system (Figure S5A). *Rspo3* overexpression phenocopied our *Sf1-Rspo1^GOF^* model in terms of sex-specific hyperplasia and ectopic proliferation at 6 weeks of age (Figure S5 B,C). However, we were not able to compare the two models directly because the expression of the *Rspo3* transgene was not uniform (Figure S5D). Thus, we conclude that overexpression of either R-spondin affects the adrenal cortex in the same manner.

### Male *Rspo1^GOF^* adrenals display abnormal macrophage accumulation

Because our transcriptomic analysis revealed an enrichment for immune-related genes among the cluster of genes specifically upregulated in male *Rspo1^GOF^*, we examined the expression of macrophage markers by immunofluorescence staining. CD68 represents a marker of inflammation abundantly expressed in macrophages and is also detected in other cell types of the myeloid lineage ^45^. CD68 is expressed in wild-type adrenals at 6 weeks of age at the border between the cortex and the medulla or the X-zone (Figure 2F). By contrast, CD68 is marking small cells scattered around the cortex in the hyperplastic *Sf1-Rspo1^GOF^* female adrenals, while in male *Sf1-Rspo1^GOF^* adrenals, CD68 marks cells increasing in size that adopt a ‘foamy’ morphology, fuse and contribute to the formation of the degenerative lesions (Figure 2F). Several genes expressed in macrophages and monocytes, as well as pan immune cell markers, are upregulated in male *Sf1-Rspo1^GOF^* adrenals (Figure S6A). Moreover, IBA1, a macrophage marker that has been shown to be induced in *Star* knockout animals^46^, was strongly expressed in male *Sf1-Rspo1^GOF^* adrenals, but not in their female counterparts (Figure S6B). Overall, our data suggest that male *Sf1-Rspo1^GOF^* adrenals develop a sex-specific inflammatory profile, characterised by increased presence of monocytes and macrophages, that culminates in cortical thinning with increased age.

### Sex specific pattern of canonical WNT signalling activation in *Rspo1^GOF^* adrenals

In order to identify the cause of sexual dimorphism in our model, we tested whether sex influences canonical WNT signalling activation. To focus on primary events, we chose an early timepoint (4 weeks), when sex specific differences in proliferation start to emerge, but degeneration is not yet obvious in male adrenals. In agreement with previous research ^15^, β-catenin shows strong membrane immunoreactivity in the zG but is absent in the inner cortex of control animals (apart from small spindle-like nuclei which do not have the characteristic morphology of steroidogenic cells). However, in *Sf1-Rspo1^GOF^* adrenals, we observed increased membrane, nuclear and perinuclear β-catenin immunoreactivity in the inner cortex, in addition to its characteristic zG pattern (Figure 3A). To quantify this disruption of the characteristic WNT signalling gradient, we performed RNA Scope *in situ* hybridisation (ISH) analysis for *Axin2* and *Wnt4*, two markers of canonical WNT signalling activation. While *Axin2* is a well-characterised target of canonical WNT signalling ^47^, *Wnt4* is thought to be a driver of canonical WNT signalling in the adrenal cortex ^21,22^. Consistent with previous data ^22^, the expression of these two genes follows a gradient of diminishing expression from the outer to the inner cortex in control animals, which was disrupted in our GOF model (Figure 3B, C). Interestingly, the expression of these two genes became sexually dimorphic in *Sf1-Rspo1^GOF^* adrenals, with *Axin2* expression being increased in males (reaching statistical significance in the outer cortex) and *Wnt4* expression being increased in females (reaching statistical significance in the inner cortex) (Figure 3D, E).

**Figure 3.**
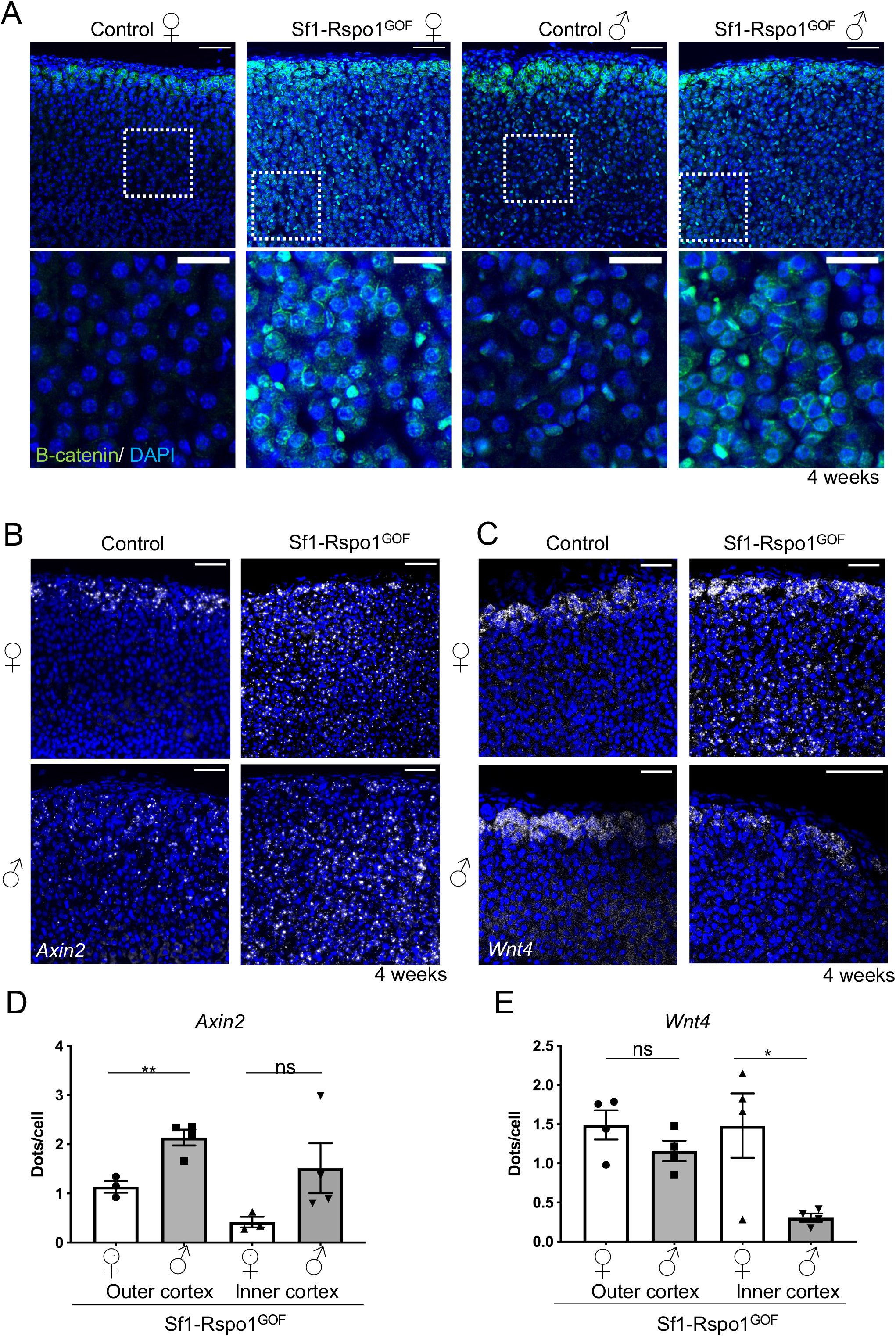
RSPO1 overexpression induces the expansion of canonical WNT activity zone in the adrenal cortex in a sexually dimorphic manner. **A**) Immunofluorescence staining for β-catenin using adrenal sections from 4 week-old mice. Scale bar: 50 μm μm (top panel), 20 μm (bottom panel). **B, C**) In situ hybridisation using the RNA Scope technology for *Axin2* (**B**), a target of canonical WNT signalling, and *Wnt4* (**C**) at 4 weeks of age. Scale bars: 50 μm. **D, E**) Graphs showing the mean number of *Axin2* (**D**) and *Wnt4* **(E**) transcripts per cell in the outer cortex or inner cortex, based on RNA Scope-based detection on adrenal sections from 4 week-old mice (error bars represent SEM, while each dot represents an independent biological replicate). Statistical analysis was performed using unpaired t-test. P value comparing male vs female outer cortex for *Axin2* P=0.0058. P value comparing male vs female inner cortex P=0.1302. P value comparing male vs female outer cortex for *Wnt4* P=0.1963. P-value comparing male vs female inner cortex P=0.0301.

### Androgen receptor signalling determines sexual dimorphism in *Rspo1^GOF^* adrenals

To explain sexual dimorphism in adrenocortical hyperplasia, it is essential to dissect the role of sex hormones versus the role of sex chromosomes. We took advantage of a sex reversal model where constitutive ectopic expression of a *Wt1-Sox9* gene in XX gonads leads to the development of testes ^48^ (Figure 4A). Sex-reversed *Sf1-Rspo1^GOF^* males (*Gt(Rosa)26Sor^cCAG-Rspo1/+^*; *Sf1-cre^Tg/0^; Wt1-Sox9^Tg/0^* XX) displayed significantly lower normalised adrenal weight than *Sf1-Rspo1^GOF^* females (*Gt(Rosa)26Sor^cCAG-Rspo1/+^*; *Sf1-cre^Tg/0^* XX) (Figure 4B) and developed vacuolated cells in the inner cortex at 6 weeks of age (Figure 4C) thus phenocopying XY *Sf1-Rspo1^GOF^* male adrenals. These data indicate that gonadal rather than chromosomal sex is responsible for the sexual dimorphism.

**Figure 4.**
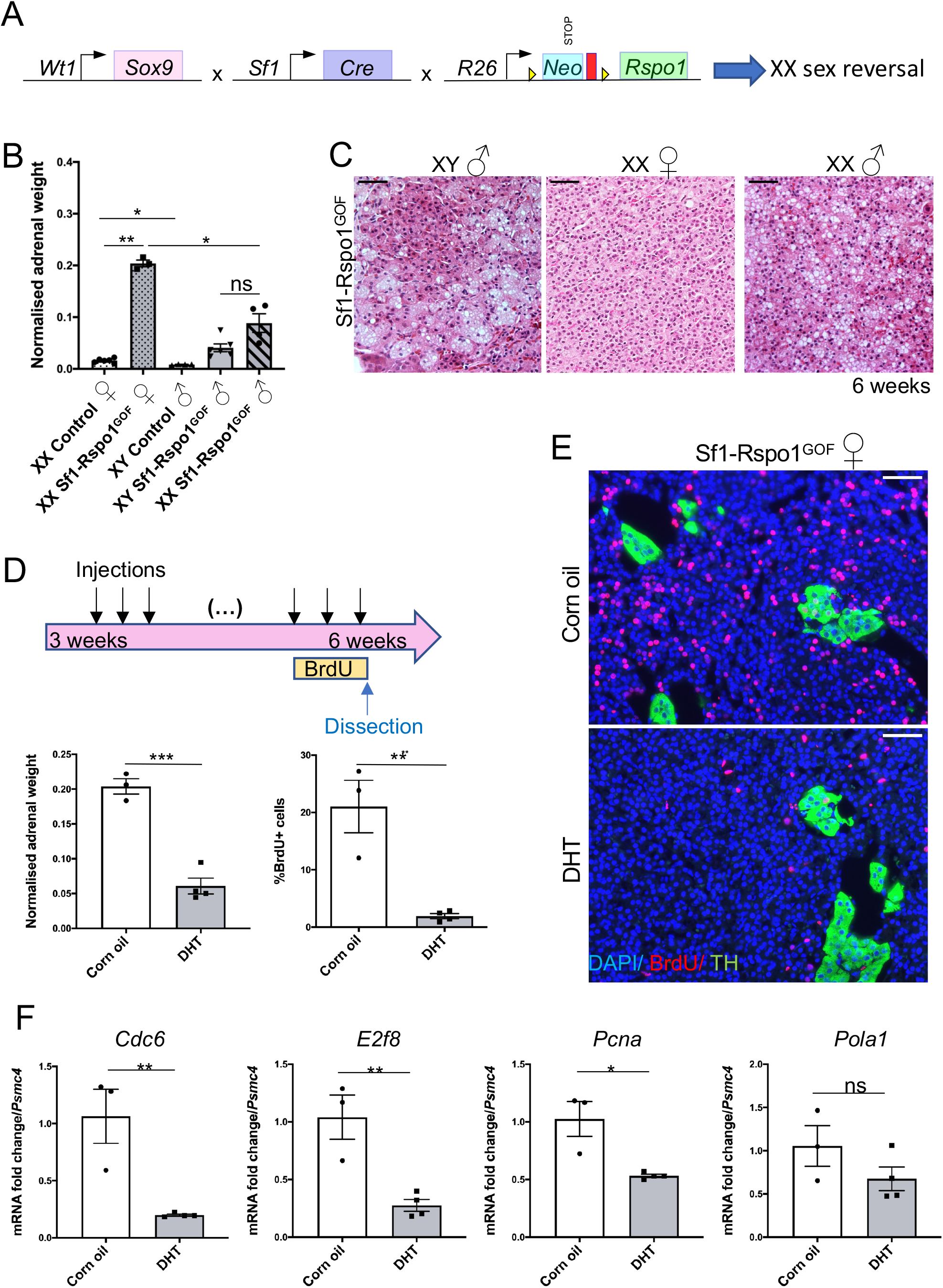
Sexually dimorphic response to RSPO1 overexpression is caused by androgens. **A**) Schematic representation of the genetic strategy to combine RSPO1 overexpression in steroidogenic tissues with female-to-male sex-reversal caused by ectopic SOX9 expression under control of a *Wt1* regulatory sequence. **B**) Mean adrenal weight normalised to whole body weight at 6 weeks. “XX” and “XY” denote sex chromosomes and “XX ♂” indicate sex-reversed Tg(Wt1-Sox9)^Tg/0^ mice. Error bars represent SEM. Statistical analysis was performed with one-way Welch’s ANOVA followed by Dunnett’s T3 multiple comparisons test. Adjusted P-values: XX Control F vs XX *Sf1-Rspo1^GOF^* F: P=0.0048, XX Control F vs XY Control M: P=0.0367, XX *Sf1-Rspo1^GOF^* F vs XX *Sf1-Rspo1^GOF^* M: P=0.0231. **C**) H&E staining on adrenal sections from 6 weeks -old XY and XX *Sf1-Rspo1^GOF^* male mice. Black arrows point to vacuolated cells that form degenerative lesions. Scale bar: 50 μm. **D**) Female *Sf1-Rspo1^GOF^* mice were treated with DHT or corn oil daily for 3 weeks during puberty. Graphs represent mean adrenal weight normalised to total body weight (top) and percentage of proliferating cells (Brdu+) in the adrenal cortex (bottom). Error bars represent SEM. Statistical analysis was conducted using unpaired two-tailed t-test (P values: P=0.0003-top graph, P=0.0043-bottom graph). **E**) Representative immunofluorescence images for BrdU and TH (marker of the medulla). Scale bar: 50 μm. **F**) RT-qPCR analysis of the expression of genes related to G1/S cell cycle transition. Graphs represent mean fold expression change comparing corn oil to DHT treated adrenals (normalised to *Psmc4* expression). Error bars represent SEM. Statistical analysis was conducted with unpaired t-test. P values: 0.1964 (*Pola1*), 0.0066 (*E2f8*), 0.0073 (*Cdc6*), 0.0117 (*Pcna*). DHT: dihydrotestosterone.

We have shown previously that androgens can suppress progenitor proliferation during normal adrenal cortex homeostasis ^33^. To test whether androgens would also modify female specific hyperplasia in our mouse model, we chose to treat *Sf1-Rspo1^GOF^* female mice daily with the androgen dihydrotestosterone (DHT) during puberty (3 to 6 weeks of age) (Figure 4D). Expression analysis revealed a slight reduction of the androgen receptor gene (*Ar*), as well as a dramatic increase in the expression of *Susd3*, a putative androgen-responsive gene in the adrenal cortex (as suggested by our RNA sequencing data and previous transcriptomic analyses ^49^) (Figure S7B). Moreover, DHT treatment dramatically reduced normalised adrenal weight and proliferation levels in the adrenal cortex (Figure 4D,E) but did not lead to histological changes such as cytoplasmic vacuolisation (Figure S7A). The reduction of proliferation levels in DHT-treated animals was accompanied by a reduction in expression levels of genes associated with G1/S transition of the cell cycle (*E2f8, Pcna, Cdc6, Pola1*) ^50^ (Figure 4F). Further, DHT treatment led to an increase in *Axin2* expression levels and to a decrease in *Wnt4* expression levels (Figure S7B), similarly to what RNA Scope ISH analysis suggested regarding the effect of sex on the expression of WNT-related genes (Figure 3C, E). Adrenal cortex homeostasis depends on complex endocrine interactions with the pituitary-hypothalamic axis and the gonads ^51^. Having established a role for DHT in suppressing hyperplasia, we asked whether this effect is direct or involves feedback loops via other organs. To answer this question, we took advantage of a conditional knockout allele for androgen receptor (*Ar*) ^52^ that we activated simultaneously with *Rspo1* transgene. To exclude extra-adrenal effects we employed the adrenal cortex-specific aldosterone synthase (AS) *Cyp11b2-Cre* line (Figure 5A) that becomes activated in the zG at birth and – due to the centripetal displacement of cortical cells – leads to recombination throughout the cortex by approximately 5 weeks. Given the slower recombination rate compared to *Sf1-Cre*, we analysed adrenals at 20 weeks of age. Similarly to *Sf1-Cre, Cyp11b2-Cre* activation of *Rspo1(AS-Rspo1^GOF^*) resulted in female adrenal hyperplasia whereas growth of the male adrenals remained comparable to wildtype controls (Figure 5B). Strikingly, simultaneous deletion of the *Ar* allele caused a significant increase of male adrenals to a weight comparable to that found in female counterparts (Figure 5B). Moreover, male *AS-Rspo1^GOF^*/*Ar KO* and female *AS-Rspo1^GOF^* adrenals appeared similar on the histological level, characterised by cortical hyperplasia (Figure S8A). BrdU-based analysis confirmed an increase in proliferation in male *Rspo1^GOF^/Ar KO* adrenals, particularly in the inner cortex. The observed variability among different animals (Figure 5C,D), is likely due to incomplete deletion of the *Ar* allele in clusters of adreno-cortical cells (Figure S8B). Taken together, our results demonstrates that androgens act directly on adrenocortical cells by engaging their cognate receptor and cause proliferation arrest, contributing to a differential susceptibility to adrenocortical hyperplasia among the sexes (Figure 6).

**Figure 5.**
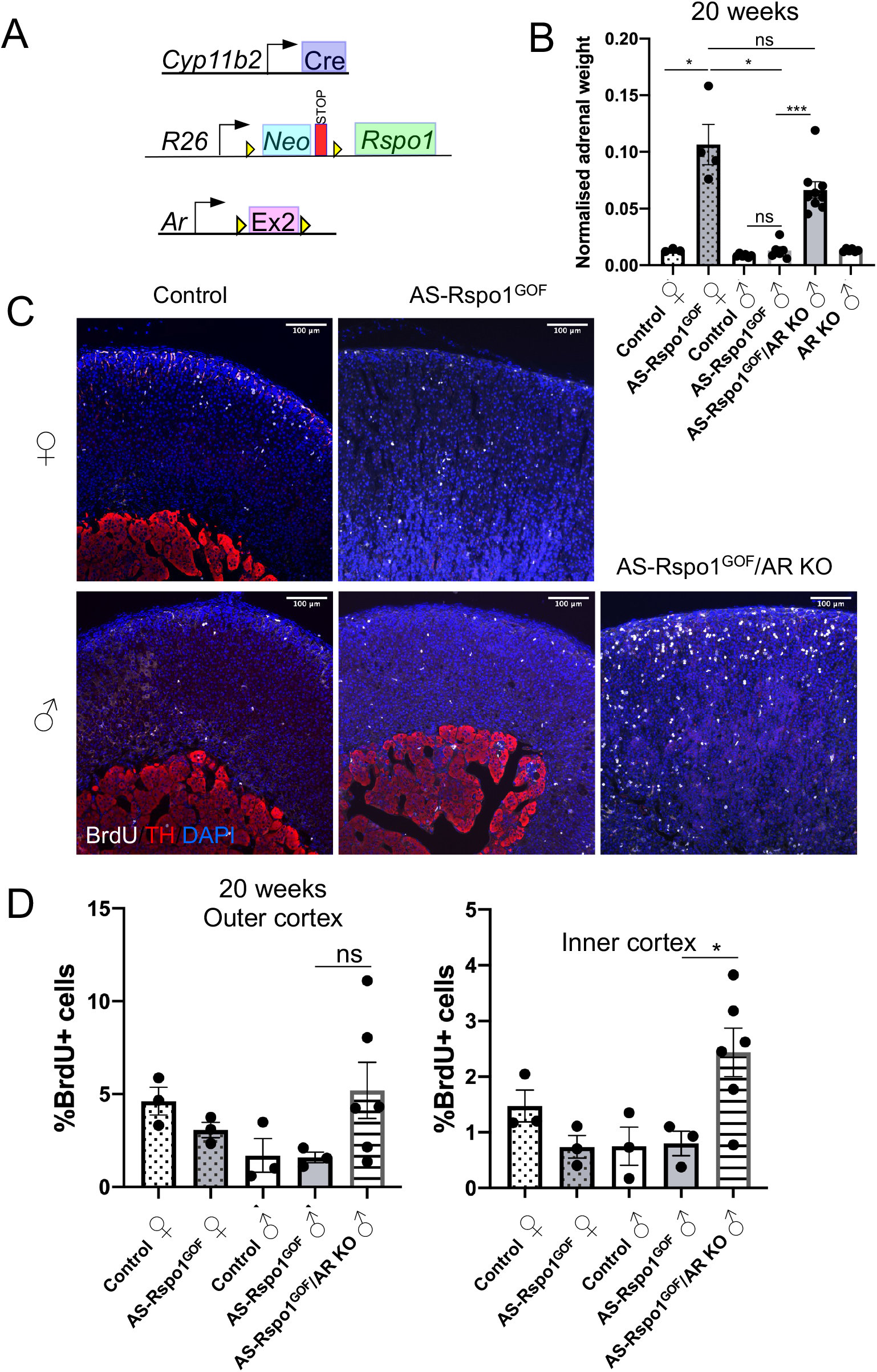
Androgen receptor deletion in male AS-Rspo1 adrenocortical cells abolishes sexual dimorphism in the development of hyperplasia. **A**) Schematic representation of the genetic strategy to knock-out AR and simultaneously overexpress RSPO1 in adrenocortical cells using the *Cyp11b2-Cre* driver. **B**) Adrenal weight normalised to body weight at 20 weeks (error bars represent SEM). Statistical analysis was performed using Welch’s one-way ANOVA followed by Dunnett’s T3 post-hoc test. Adjusted P values: Control F vs *AS-Rspo1^GOF^* F: P=0.0454, Control M vs *AS-Rspo1^GOF^* M: P=0.5777, *AS-Rspo1^GOF^* F vs *AS-Rspo1^GOF^* M: P=0.0468, *AS-Rspo1^GOF^* F vs *AS-Rspo1^GOF^/Ar KO* M: P=0.3370, *AS-Rspo1^GOF^* M vs *AS-Rspo1^GOF^/Ar KO* M: O=0.0001. **C**) Representative immunofluorescence images for Tyrosine hydroxylase (TH) and BrdU, indicating proliferating cells. Scale bar: 100 μm. **D**) BrdU proliferation analysis shown as percentage of proliferating cells over total number of cells in the adrenal cortex of 20 week-old mice (error bars represent SEM). The area close to the capsule (outer cortex) is distinguished from the deeper layers (inner cortex). Unpaired t-test was employed to compare values for *AS-Rspo1^GOF^* male and *AS-Rspo1^GOF^/Ar KO* male. Adjusted p-values: 0.1478 (outer cortex), 0.0410 (inner cortex).

**Figure 6.**
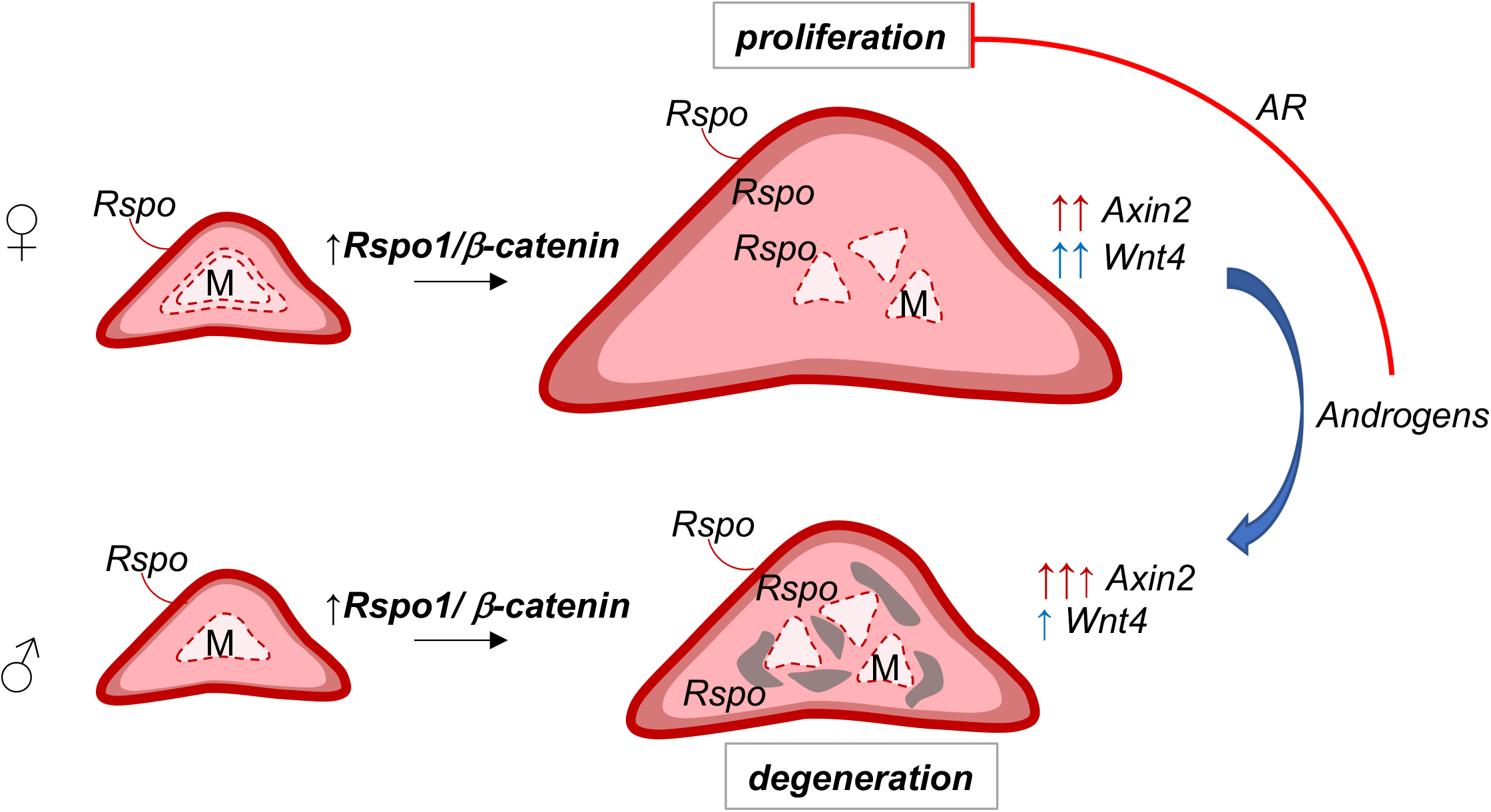
Sex-specific effects of *Rspo1* overexpression in the adrenal cortex. Localised expression of R-spondins (by the capsule) maintain high levels of WNT/β-catenin signalling and proliferation in the outer cortex. Ectopic expression of *Rspo1* by steroidogenic cells leads to increased WNT/β-catenin signalling in the inner cortex, as indicated by ectopic expression of *Wnt4* and *Axin2*, and ectopic proliferation that causes adrenocortical hyperplasia and fragmentation of the medulla. On the contrary, in male adrenals or in female adrenals treated with androgens (DHT), ectopic *Rspo1* causes differential effects on *Wnt4* and *Axin2* expression, while ectopic proliferation is prevented via activation of AR signalling.

## Discussion

Canonical WNT/β-catenin signalling gradients determine a plethora of processes, ranging from embryonic development to the maintenance of adult stem cell niches, while their dysregulation is linked to carcinogenesis in humans ^20^. High β-catenin levels in the zG of the mouse adrenal cortex, supported by secreted R-spondins, are paramount to the maintenance of regular renewal and zonation ^21,53^. Failure to turn-off β-catenin signalling in the zF due to the overexpression of a constitutively active form of the protein impedes transdifferentiation, leading to ectopic zG expansion ^16,17^. By contrast, *Rspo1* overexpression (as shown in this study) as well as *Znrf3* deletion ^22^ leads to ectopic proliferation, but not an expansion of the zG despite the presence of *Wnt4* and *Rspo1* in deeper zones of *Rspo1^GOF^* adrenals. These observations suggests that additional factors - perhaps released from the capsule - are required to ensure strong β-catenin activity in subcapsular cells. Interesting candidates for such molecules are fibroblast growth factors (FGF) that have been recently shown to be required for normal zG formation ^15^.

We found a striking, androgen-dependent, sexual dimorphism in phenotypic development in *Rspo1^GOF^* mice with two major pillars of tumourigenesis regulated in a sex hormone-specific manner. Firstly, recruitment of macrophages, monocytes and dendritic cells leaves a more prominent gene expression signature in males and manifests as foamy cell formation, cytoplasmic vacuolisation and tissue degeneration at the histological level. On the other hand, hyperproliferation is female-specific, suppressed by androgen administration and AR signalling, even though these manipulations do not directly cause foamy cell formation and degeneration at the histological level. Hence, we were able to analyse proliferation arrest as an androgen-dependent phenomenon, separate from endocrine manifestations and hypercortisolism, acknowledging that the immune cell activation aspect deserves further investigation in a separate study.

A high proliferation index is an important feature of ACC, associated with malignant rather than benign tumours and with worse prognosis ^54^. Moreover, dysregulation of WNT/β-catenin signalling and cell cycle regulation (via the p53/Retinoblastoma-associated protein (RB1) axis) are significant hallmarks of ACC and the pathways most frequently affected by driver mutations ^24,25^. In this context, the findings of the present study can contribute to our greater understanding of sex bias in ACC frequency and the development of personalised therapies. Importantly, we show that ectopic expression of *Rspo1* alone is however insufficient to cause ACC in mice, and only 1/8 females and 0/8 males showed carcinoma at 12 months. This is in agreement with a recent report that demonstrated that additional alterations such as *Tp53* deletion are required to induce aggressive tumours ^55^. In future studies it will be important to test whether androgen administration in the context of such ACC models also suppresses proliferation and tumour growth. Of note, it has been described that DHT treatment of human ACC cells leads to growth arrest ^56^.

Androgens are potent suppressors of the HPA axis at the level of the hypothalamus and the pituitary gland, which in turn regulate corticosterone production by the adrenal cortex ^57^. Distinguishing direct versus indirect effects of sex hormones in gonadectomy and/or DHT treatment experiments has therefore been difficult. We show here that removing the cognate DHT receptor from adrenocortical cells (*Ar* deletion) renders male adrenals susceptible to hyperplasia and hyperproliferation and abolishes sexual dimorphism. Thus, this is the first report of AR signalling directly suppressing proliferation in the adrenal cortex. Of note, it has also been shown before that androgen receptor signalling influences adrenal weight and X-zone regression ^58^. On the molecular, it is possible that AR directly antagonises WNT/β-catenin signalling, as it has been suggested before for epidermal stem cells^59^. As another example, in prostate cancer cells, AR interacts with β-catenin and has been suggested to negatively regulate its transcriptional activity by competing with TCF for binding ^60^, although the two pathways synergistically promote prostate tumorigenesis ^61^. Despite these associations, we did not find evidence for a sex-specific global reduction of β-catenin signalling, but instead an opposing effect of sex on the expression of two canonical WNT-related genes, *Axin2* and *Wnt4*. Another possibility would be that AR directly suppresses transcription of cell cycle genes independently of WNT signalling. Interestingly, it has already been reported that AR acts as a transcriptional repressor for a subset of DNA replication genes by recruiting RB1 to their promoters ^62^.

Our report of AR being a suppressor of proliferation in the adrenal cortex might be surprising, given the well-characterised association of AR not only with prostate tumorigenesis, but also with promoting normal prostate growth and benign prostate hyperplasia ^63^. However, even in the context of prostate cancer, AR can have opposing effects on cell cycle regulation, depending on ligand concentration, duration of treatment and the extent of gain-of-function alterations in AR signalling in the context of malignant transformation ^62,64,65^. Moreover, it has been suggested that AR acts as a tumour suppressor in the context of breast cancer ^66^. Our study contributes to the understanding of AR signalling complexity and tissue specificity and highlights one prominent cause of sexual dimorphism in a non-reproductive organ. At the same, our findings about the role of circulating androgens in adrenal hyperplasia can explain sex bias in adrenal disease and aid the development of future personalised therapies.

## Materials and Methods

### Animal husbandry and genetics

All animal work was conducted according to national and international guidelines and approved by the local ethical committee (CIEPAL: APAFIS#6001-201606281711255 v6, APAFIS#14137-2018030216239792 v1) and the French ministry for agriculture. The mouse strains used in this study have been reported previously: *Rspo1*^*GOF* 35,36^, *Sf1-Cre*^37^, *Wt1-Sox9* ^48^, *Rspo3^GOF^* ^44^, *Cyp11b2-Cre^13^, Ar^ffox^* ^52^. Mice heterozygous for the *Rspo1^GOF^* allele and the Sf1-Cre are referred to as *“Sf1-Rspo1^GOF^” (Gt(Rosa)26Sor^cCAG-Rspo1/+^*; *Sf1-Cre*^Tg/0^), while mice with genotypes that do not permit the expression of the GOF allele are referred to as “controls” (*Gt(Rosa)26Sor^+/+^*; *Sf1-Cre*^Tg/0^, *Gt(Rosa)26Sor^+/+^*; *Sf1-Cre* or *Gt(Rosa)26Sor^cCAG-Rspo1/+^*; *Sf1-Cre*^0/0^). The expression of a *Wt1-Sox9* allele distinguishes sex-reversed *Sf1-Rspo1^GOF^* males (*Gt(Rosa)26Sor^cCAG-Rspo1/+^*; *Sf1-Cre*^Tg/0^; *Wt1-Sox9*^Tg/0^ XX) from *Sf1-Rspo1^GOF^* females (*Gt(Rosa)26Sor^cCAG-Rspo1/+^*; *Sf1-Cre*^Tg/0^; *Wt1-Sox9*^0/0^ XX) and males (*Gt(Rosa)26Sor^cCAG-Rspo1/+^*; *Sf1-Cre*^Tg/0^; *Wt1-Sox9*^0/0^ XY). Mice heterozygous for the *Rspo1^GOF^* allele and the *Cyp11b2* (aldosterone synthase)-Cre are referred to as *‘AS-Rspo1^GOF^’ (Gt(Rosa)26Sor^cCAG-Rspo1/+^*; *Cyp11b2^Cre/+^*), compared to appropriate controls as described above for the Sf1-Cre, and *‘AS-Rspo1^GOF^/Ar KO’* when the gene encoding for androgen receptor is deleted in males (*Gt(Rosa)26Sor^cCAG-Rspo1/+^*; *Cyp11b2^Cre/+^*; *Ar^flox/Y^*). Mouse lines were maintained on a mixed genetic background. Both males and females were analysed at various ages as indicated in the main text, while littermates were preferentially compared.

### Immunofluorescence and histology

For immunofluorescence (IF) and H&E analysis of paraffin-embedded samples, mouse left adrenal tissues were fixed overnight at 4°C with 4% paraformaldehyde, progressively dehydrated and paraffin embedded. For H&E staining, sections of 5 mm were rehydrated and stained with eosin and Mayer’s hematoxylin (3 minutes each); before being dehydrated again and mounted using an anhydrous mounting medium. Carcinoma and adenoma formation was verified by expert pathologists. For IF, 5 mm sections were unmasked with PT Link (Dako Agilent Pathology Solutions) at pH 6 or 9. Sections were blocked for 1h with 10% Normal Donkey Serum (Jackson Immunoresearch), 3% BSA (Sigma) and 0.1% Tween-20 (Sigma). Primary antibodies were applied overnight at 4°C diluted in 3% NDS, 3% BSA, 0.1% Tween-20 as explained in Supplementary table S2. The next day, sections were incubated at room temperature with secondary antibodies diluted 1:400 in PBS and mounted in antifade mounting medium with DAPI (Vectashield). The Mouse-on-Mouse immunodetection kit (Vector Laboratories) was used to improve the signal for mouse primary antibodies according to manufacturer’s instructions. For anti-AR and IBA1 immunohistochemistry, sections were incubated with biotinylated secondary antibodies and the signal was revealed using the Vectastain Elite ABC-HRP peroxidase kit and the ImmPACT NovaRED HRP substrate (Vector laboratories). Counterstaining was performed using 50% Harris hematoxylin for 30 seconds followed by incubation in 0.1% sodium bicarbonate solution for 1 minute at room temperature.

### Proliferation quantification

For proliferation quantification in the adrenal cortex, 1 mg/ml of BrdU (Sigma) was dissolved in autoclaved water with 2% sugar and given to mice as drinking water for 3 days. Anti-BrdU immunofluorescence was conducted as described above. Quantification was performed using the HALO image analysis platform (Indica Labs) on whole section mosaic images obtained with the Vectra Polaris imaging system (Akoya Biosciences), excluding the medulla based on TH staining. The number of BrdU-positive cells was expressed as percentage of the total number of cells in each zone based on DAPI staining. When the distinction between ‘outer’ and ‘inner’ cortex is made, we refer to the zone <80 mm from the capsule and >80 mm from the capsule, respectively. At least three biological replicates (individual mice) and three non-consecutive sections for each biological replicate were analysed.

### RNA Scope ISH

Adrenal sections were fixed with 4% paraformaldehyde overnight at room temperature and paraffin embedded. Fresh 5 μm sections were subjected to single-molecule ISH using the RNA Scope 2.5 High Definition-Red assay (ACD Biotechne) according to manufacturer’s instructions. Images were acquired with a Zeiss apotome upright microscope or a Zeiss LSM NLO 780 confocal microscope. For quantification purposes, whole-section mosaics images were acquired with the Vectra Polaris imaging system (Akoya Biosciences) and quantified using the FISH Multiplex v1.1 module of the HALO image analysis platform (Indica Labs). When the distinction between ‘outer’ and ‘inner’ cortex is made, we refer to the zone <80 mm from the capsule and >80 mm from the capsule, respectively.

### Gene expression analysis

RNA was extracted from mouse right adrenals using the RNeasy mini kit (Qiagen) according to the manufacturer’s instructions. cDNA synthesis was performed using M-MLV reverse transcriptase (Invitrogen) and random primers. The obtained cDNA was used as template in a real-time quantitative PCR reaction using the SYBR Green master mix (Roche) and a LightCycler 1.5 (Roche) or a QuantStudio 5 thermocycler (Applied Biosystems). Expression levels were normalised to *Psmc4* housekeeping gene, analysed using the 2^-ddCt^ method and presented as fold-change values compared to a reference sample. The primers used are listed in supplementary table 1.

### Hormonal treatment and analysis of hormone levels in plasma

*Sf1-Rspo1^GOF^* females were injected twice daily subcutaneously with 5a-androstan-17b-ol-3-one (Sigma) (37.5 ug in 5% ethanol and corn oil) or oil only from 3 to 6 weeks of age, when they were sacrificed.

For the measurement of adrenal steroids in mouse plasma, 6-week-old animals were sacrificed in the morning and core trunk blood was collected in tubes containing 5 μl of 0.5M EDTA. The samples were centrifuged for 5 minutes at 4000 g at 4°C to separate the plasma, which was promptly frozen at −80°C until analysis. Steroids hormones were quantified by LC-MS/MS as described previously^67^.

### RNA Sequencing and analysis

RNA was extracted from mouse right adrenals using the RNeasy mini kit (Qiagen) according to the manufacturer’s instructions. Four biological replicates were analysed for each group (control male, control female, *Sf1-Rspo1^GOF^* male, *Sf1-Rspo1^GOF^* female). The sample’s quality was assessed using a Bioanalyzer 2100 (Agilent technologies) and a RIN cut-off value of 7.0 was applied.

Library preparation and sequencing, as well as differential expression analysis were conducted by Novogene Co. Briefly, library preparation was conducted using a NEBNext Ultra RNA Library Prep kit for Illumina (NEB). After cluster generation, the library preparations were sequenced on an Illumina platform and 125/150 bp paired-end reads were generated. Raw data (raw reads) of fastq format were firstly processed through in-house perl scripts. In this step, clean data (clean reads) were obtained by removing reads containing adapter, reads containing poly-N and low quality reads from raw data. At the same time, Q20, Q30 and GC content the clean data were calculated. All the downstream analyses were based on the clean data with high quality. Reference genome and gene model annotation files were downloaded from genome website directly. Index of the reference genome was built using Bowtie v2.2.3 and paired-end clean reads were aligned to the reference genome using TopHat v2.0.12. HTSeq v0.6.1 was used to count the reads numbers mapped to each gene. FPKM of each gene was calculated based on the length of the gene and reads count mapped to this gene. Differential expression analysis was performed using the DESeq R package (1.18.0).

The resulting P-values were adjusted using the Benjamini and Hochberg’s approach for controlling the false discovery rate. Genes with an adjusted P-value < 0.05 found by DESeq were assigned as differentially expressed.

For downstream analyses, principal component analysis (PCA) plot and heatmaps were designed using the Phantasus website tools^68^, using log10(FPKM+1) expression values from differentially expressed genes as template. Enriched gene sets were calculated using the Molecular Signatures database (Broad Institute) with an FDR q-value threshold of 0.05. GSEA analysis^69^ was conducted after DESeq2 was used via the GenePattern platform^70^ to calculate differentially expressed genes between male and female GOF adrenals.

### Statistical analysis

Statistical analysis was conducted as indicated in each figure legend using the GraphPad Prism 7 software.

## Acknowledgments

We would like to thank the staff of the animal facility for their dedication and S. Rekima (histology platform) for help in analysing RNA-Scope quantification. We are grateful to S. Sacco, V. Vidal and R. Bandiera for initial analysis of the *Rspo1^GOF^* phenotype, to D. Breault (Harvard, USA) for the *AS-cre*, K. Parker for the *Sf1-Cre* and AR and J. Hilkins for the *Rspo3^GOF^* alleles.

## Funding

La Ligue Contre le Cancer: Equipe Labelisée 2018 (AS)

Agence National de la Recherche : ANR-11-LABX-0028-01, ANR-18-CE14-0012 (AS) Worldwide Cancer Research foundation : WWCR 18-0437 (AS). R.L. was supported by an FRM Fondation de la Recherche Medicale: FRM SPF201809007141 (RL).

Deutsche Forschungsgemeinschaft: CRC/Transregio 205/1, Project No. 314061271 (AS, MP, NB)

## Author contributions

RL, AG, AS designed the work. RL, AG, AT performed all experiments if not otherwise stated. MP and NB performed steroidomics analysis. SY and AdB performed histological analysis of aged animals. ERMB and FC provided mouse lines. RL and AS wrote the manuscript. All authors read the manuscript and provided input.

## Data and materials availability

RNA-Seq data can be accessed with the GEO Series accession number GSE178958. All other data are available in the main text or the supplementary materials.

## References

1 Karp NA, Mason J, Beaudet AL, Benjamini Y, Bower L, Braun RE et al. Prevalence of sexual dimorphism in mammalian phenotypic traits. Nat Commun 2017; 8: 15475.

2 Clocchiatti A, Cora E, Zhang Y, Dotto GP. Sexual dimorphism in cancer. Nat Rev Cancer 2016; 16: 330–339.

3 Klein SL, Flanagan KL. Sex differences in immune responses. Nat Rev Immunol 2016; 16: 626–638.

4 Yu X, Li S, Xu Y, Zhang Y, Ma W, Liang C et al. Androgen Maintains Intestinal Homeostasis by Inhibiting BMP Signaling via Intestinal Stromal Cells. Stem Cell Reports 2020; 0. doi:10.1016/j.stemcr.2020.08.001.

5 Chen X, Mcclusky R, Chen J, Beaven SW, Tontonoz P. The Number of X Chromosomes Causes Sex Differences in Adiposity in Mice. PLoS Genet 2012; 8: 1002709.

6 Pignatti E, Leng S, Carlone DL, Breault DT. Regulation of zonation and homeostasis in the adrenal cortex. Mol Cell Endocrinol 2017; 441: 146–155.

7 Huang C-CJ, Kang Y. The transient cortical zone in the adrenal gland: the mystery of the adrenal X-zone. J Endocrinol 2019; 241: R51–R63.

8 Lyraki R, Schedl A. Adrenal cortex renewal in health and disease. Nat Rev Endocrinol 2021;:1–14.

9 King P, Paul A, Laufer E. Shh signaling regulates adrenocortical development and identifies progenitors of steroidogenic lineages. Proc Natl Acad Sci U S A 2009; 106: 21185–21190.

10 Ching S, Vilain E. Targeted disruption of Sonic Hedgehog in the mouse adrenal leads to adrenocortical hypoplasia. genesis 2009; 47: 628–637.

11 Finco I, Lerario AM, Hammer GD. Sonic Hedgehog and WNT Signaling Promote Adrenal Gland Regeneration in Male Mice. Endocrinology 2018; 159: 579–596.

12 Chang SP, Morrison HD, Nilsson F, Kenyon CJ, West JD, Morley SD. Cell proliferation, movement and differentiation during maintenance of the adult mouse adrenal cortex. PLoS One 2013; 8. doi:10.1371/journal.pone.0081865.

13 Freedman BD, Kempna PB, Carlone DL, Shah M, Guagliardo NA, Barrett PQ et al. Adrenocortical zonation results from lineage conversion of differentiated zona glomerulosa cells. Dev Cell 2013; 26: 666–673.

14 Kim AC, Reuter AL, Zubair M, Else T, Serecky K, Bingham NC et al. Targeted disruption of beta-catenin in Sf1-expressing cells impairs development and maintenance of the adrenal cortex. Development 2008; 135: 2593–602.

15 Leng S, Pignatti E, Khetani RS, Shah MS, Xu S, Miao J et al. β-Catenin and FGFR2 regulate postnatal rosette-based adrenocortical morphogenesis. Nat Commun 2020; 11. doi:10.1038/s41467-020-15332-7.

16 Berthon A, Sahut-Barnola I, Lambert-Langlais S, de Joussineau C, Damon-Soubeyrand C, Louiset E et al. Constitutive β-catenin activation induces adrenal hyperplasia and promotes adrenal cancer development. Hum Mol Genet 2010; 19: 1561–1576.

17 Pignatti E, Leng S, Yuchi Y, Borges KS, Guagliardo NA, Shah MS et al. Beta-Catenin Causes Adrenal Hyperplasia by Blocking Zonal Transdifferentiation. Cell Rep 2020; 31. doi:10.1016/j.celrep.2020.107524.

18 Hao HX, Xie Y, Zhang Y, Zhang O, Oster E, Avello M et al. ZNRF3 promotes Wnt receptor turnover in an R-spondin-sensitive manner. Nature 2012; 485: 195–202.

19 Zebisch M, Xu Y, Krastev C, Macdonald BT, Chen M, Gilbert RJC et al. Structural and molecular basis of ZNRF3/RNF43 transmembrane ubiquitin ligase inhibition by the Wnt agonist R-spondin. Nat Commun 2013; 4. doi:10.1038/ncomms3787.

20 Nusse R, Clevers H. Wnt/ß-Catenin Signaling, Disease, and Emerging Therapeutic Modalities. Cell 2017; 169: 985–999.

21 Vidal V, Sacco S, Rocha AS, Da Silva F, Panzolini C, Dumontet T et al. The adrenal capsule is a signaling center controlling cell renewal and zonation through Rspo3. Genes Dev 2016; 30: 1389–1394.

22 Basham KJ, Rodriguez S, Turcu AF, Lerario AM, Logan CY, Rysztak MR et al. A ZNRF3-dependent Wnt/ß-catenin signaling gradient is required for adrenal homeostasis. Genes Dev 2019; 33: 209–220.

23 Heikkilä M, Peltoketo H, Leppäluoto J, Ilves M, Vuolteenaho O, Vainio S. Wnt-4 Deficiency Alters Mouse Adrenal Cortex Function, Reducing Aldosterone Production. Endocrinology 2002; 143: 4358–4365.

24 Assié G, Letouzé E, Fassnacht M, Jouinot A, Luscap W, Barreau O et al. Integrated genomic characterization of adrenocortical carcinoma. Nat Genet 2014; 46: 607–612.

25 Zheng S, Cherniack AD, Dewal N, Moffitt RA, Danilova L, Murray BA et al. Comprehensive pan-genomic characterization of adrenocortical carcinoma. Cancer Cell 2016; 29: 723–736.

26 Lyraki R, Schedl A. The sexually dimorphic adrenal cortex: Implications for adrenal disease. Int J Mol Sci 2021; 22.

27 Lacroix A, Feelders RA, Stratakis CA, Nieman LK. Cushing’s syndrome. Lancet. 2015; 386: 913–927.

28 Lindholm J, Juul S, Jørgensen JOL, Astrup J, Bjerre P, Feldt-Rasmussen U et al. Incidence and Late Prognosis of Cushing’s Syndrome: A Population-Based Study 1. J Clin Endocrinol Metab 2001; 86: 117–123.

29 Ayala-Ramirez M, Jasim S, Feng L, Ejaz S, Deniz F, Busaidy N et al. Adrenocortical carcinoma: Clinical outcomes and prognosis of 330 patients at a tertiary care Center. Eur J Endocrinol 2013; 169: 891–899.

30 Scollo C, Russo M, Trovato MA, Sambataro D, Giuffrida D, Manusia M et al. Prognostic factors for adrenocortical carcinoma outcomes. Front Endocrinol (Lausanne) 2016; 7. doi:10.3389/fendo.2016.00099.

31 Luton JP, Cerdas S, Billaud L, Thomas G, Guilhaume B, Bertagna X et al. Clinical features of adrenocortical carcinoma, prognostic factors, and the effect of mitotane therapy. N Engl J Med 1990; 322: 1195–1201.

32 Bielohuby M, Herbach N, Wanke R, Maser-Gluth C, Beuschlein F, Wolf E et al. Growth analysis of the mouse adrenal gland from weaning to adulthood: time-and gender-dependent alterations of cell size and number in the cortical compartment. Am J Physiol Metab 2007; 293: E139–E146.

33 Grabek A, Dolfi B, Klein B, Jian-Motamedi F, Chaboissier M-C, Schedl A. The Adult Adrenal Cortex Undergoes Rapid Tissue Renewal in a Sex-Specific Manner. Cell Stem Cell 2019; 25: 290–296.e2.

34 Dumontet T, Sahut-Barnola I, Septier A, Montanier N, Plotton I, Roucher-Boulez F et al. PKA signaling drives reticularis differentiation and sexually dimorphic adrenal cortex renewal. JCI insight 2018; 3. doi:10.1172/jci.insight.98394.

35 De Cian MC, Pauper E, Bandiera R, Vidal VP, Sacco S, Gregoire EP et al. Amplification of R-spondin1 signaling induces granulosa cell fate defects and cancers in mouse adult ovary. Oncogene 2016. doi:10.1038/onc.2016.191.

36 Rocha AS, Vidal V, Mertz M, Kendall TJ, Charlet A, Okamoto H et al. The Angiocrine Factor Rspondin3 Is a Key Determinant of Liver Zonation. Cell Rep 2015; 13: 1757–1764.

37 Bingham NC, Verma-Kurvari S, Parada LF, Parker KL. Development of a steroidogenic factor 1/Cre transgenic mouse line. genesis 2006; 44: 419–424.

38 Sahut-Barnola I, Lefrancois-Martinez AM, Jean C, Veyssiere G, Martinez A. Adrenal tumorigenesis targeted by the corticotropin-regulated promoter of the aldo-keto reductase AKR1B7 gene in transgenic mice. In: Endocrine Research. Marcel Dekker Inc., 2000, pp 885–898.

39 Barrett T, Wilhite SE, Ledoux P, Evangelista C, Kim IF, Tomashevsky M et al. NCBI GEO: Archive for functional genomics data sets - Update. Nucleic Acids Res 2013; 41. doi:10.1093/nar/gks1193.

40 Fevr T, Robine S, Louvard D, Huelsken J. Wnt/ß-Catenin Is Essential for Intestinal Homeostasis and Maintenance of Intestinal Stem Cells. Mol Cell Biol 2007; 27: 7551–7559.

41 Sansom OJ, Meniel VS, Muncan V, Phesse TJ, Wilkins JA, Reed KR et al. Myc deletion rescues Apc deficiency in the small intestine. Nature 2007; 446: 676–679.

42 Xie X, Lu J, Kulbokas EJ, Golub TR, Mootha V, Lindblad-Toh K et al. Systematic discovery of regulatory motifs in human promoters and 3’ UTRs by comparison of several mammals. Nature 2005; 434: 338–345.

43 Fischer M, Grossmann P, Padi M, DeCaprio JA. Integration of TP53, DREAM, MMB-FOXM1 and RB-E2F target gene analyses identifies cell cycle gene regulatory networks. Nucleic Acids Res 2016; 44: 6070–6086.

44 Hilkens J, Timmer NC, Boer M, Ikink GJ, Schewe M, Sacchetti A et al. RSPO3 expands intestinal stem cell and niche compartments and drives tumorigenesis. Gut 2017; 66:1095–1105.

45 Chistiakov DA, Killingsworth MC, Myasoedova VA, Orekhov AN, Bobryshev Y V. CD68/macrosialin: Not just a histochemical marker. Lab Investig 2017; 97: 4–13.

46 Ishii T, Mitsui T, Suzuki S, Matsuzaki Y, Hasegawa T. A Genome-Wide Expression Profile of Adrenocortical Cells in Knockout Mice Lacking Steroidogenic Acute Regulatory Protein. Endocrinology 2012; 153: 2714–2723.

47 Jho E, Zhang T, Domon C, Joo C-K, Freund J-N, Costantini F. Wnt/ß-Catenin/Tcf Signaling Induces the Transcription of Axin2, a Negative Regulator of the Signaling Pathway. Mol Cell Biol 2002; 22: 1172–1183.

48 Vidal VPI, Chaboissier M-C, de Rooij DG, Schedl A. Sox9 induces testis development in XX transgenic mice. Nat Genet 2001; 28: 216–217.

49 El Wakil A, Mari B, Barhanin J, Lalli E. Genomic analysis of sexual dimorphism of gene expression in the mouse adrenal gland. Horm Metab Res 2013; 45: 870–873.

50 Fischer M, Müller GA. Cell cycle transcription control: DREAM/MuvB and RB-E2F complexes. Crit. Rev. Biochem. Mol. Biol. 2017; 52: 638–662.

51 Goel N, Workman JL, Lee TT, Innala L, Viau V. Sex Differences in the HPA Axis. In: Comprehensive Physiology. John Wiley & Sons, Inc.: Hoboken, NJ, USA, 2014, pp 1121–1155.

52 De Gendt K, Swinnen J V., Saunders PTK, Schoonjans L, Dewerchin M, Devos A et al. A Sertoli cell-selective knockout of the androgen receptor causes spermatogenic arrest in meiosis. Proc Natl Acad Sci U S A 2004; 101: 1327–1332.

53 Kim AC, Reuter AL, Zubair M, Else T, Serecky K, Bingham NC et al. Targeted disruption β-catenin in Sf1-expressing cells impairs development and maintenance of the adrenal cortex. Development 2008; 135: 2593–2602.

54 Crona J, Beuschlein F. Adrenocortical carcinoma — towards genomics guided clinical care. Nat. Rev. Endocrinol. 2019; 15: 548–560.

55 Borges KS, Pignatti E, Leng S, Kariyawasam D, Ruiz-Babot G, Ramalho FS et al. Wnt/ß-catenin activation cooperates with loss of p53 to cause adrenocortical carcinoma in mice. Oncogene 2020; 39: 5282–5291.

56 Rossi R, Zatelli MC, Valentini A, Cavazzini P, Fallo F, Del Senno L et al. Evidence for androgen receptor gene expression and growth inhibitory effect of dihydrotestosterone on human adrenocortical cells. J Endocrinol 1998; 159: 373–380.

57 Seale J V., Wood SA, Atkinson HC, Harbuz MS, Lightman SL. Gonadal steroid replacement reverses gonadectomy-induced changes in the corticosterone pulse profile and stress-induced hypothalamic-pituitary-adrenal axis activity of male and female rats. J Neuroendocrinol 2004; 16: 989–998.

58 Gannon A-L, O’Hara L, Mason JI, Jørgensen A, Frederiksen H, Milne L et al. Androgen receptor signalling in the male adrenal facilitates X-zone regression, cell turnover and protects against adrenal degeneration during ageing. Sci Rep 2019; 9. doi:10.1038/s41598-019-46049-3.

59 Kretzschmar K, Cottle DL, Schweiger PJ, Watt FM. The Androgen Receptor Antagonizes Wnt/ß-Catenin Signaling in Epidermal Stem Cells. J Invest Dermatol 2015; 135: 2753–2763.

60 Mulholland DJ, Read JT, Rennie PS, Cox ME, Nelson CC. Functional localization and competition between the androgen receptor and T-cell factor for nuclear β-catenin: A means for inhibition of the Tcf signaling axis. Oncogene 2003; 22: 5602–5613.

61 Lee SH, Luong R, Johnson DT, Cunha GR, Rivina L, Gonzalgo ML et al. Androgen signaling is a confounding factor for ß-catenin-mediated prostate tumorigenesis. Oncogene 2016; 35: 702–714.

62 Gao S, Gao Y, Hansen H, Chen S, Balk SP, Cai C. Androgen Receptor Tumor Suppressor Function Is Mediated by Recruitment of Retinoblastoma Protein Accession Numbers GSE76141. CellReports 2016; 17: 966–976.

63 Dai C, Heemers H, Sharifi N. Androgen signaling in prostate cancer. Cold Spring Harb Perspect Med 2017; 7. doi:10.1101/cshperspect.a030452.

64 Chatterjee P, Schweizer MT, Lucas JM, Coleman I, Nyquist MD, Frank SB et al. Supraphysiological androgens suppress prostate cancer growth through androgen receptor-mediated DNA damage. J Clin Invest 2019; 129: 4245–4260.

65 Litvinov I V., Vander Griend DJ, Antony L, Dalrymple S, De Marzo AM, Drake CG et al. Androgen receptor as a licensing factor for DNA replication in androgen-sensitive prostate cancer cells. Proc Natl Acad Sci U S A 2006; 103: 15085–15090.

66 Hickey TE, Selth LA, Chia KM, Laven-Law G, Milioli HH, Roden D et al. The androgen receptor is a tumor suppressor in estrogen receptor–positive breast cancer. Nat Med 2021; 27: 310–320.

67 Peitzsch M, Dekkers T, Haase M, Sweep FCGJ, Quack I, Antoch G et al. An LC-MS/MS method for steroid profiling during adrenal venous sampling for investigation of primary aldosteronism. J Steroid Biochem Mol Biol 2015; 145: 75–84.

68 Zenkova D., Kamenev V., Sablina R., Artyomov M. SA. Phantasus: visual and interactive gene expression analysis. https://genome.ifmo.ru/phantasus. doi:10.18129/B9.bioc.phantasus.

69 Subramanian A, Tamayo P, Mootha VK, Mukherjee S, Ebert BL, Gillette MA et al. Gene set enrichment analysis: A knowledge-based approach for interpreting genome-wide expression profiles. Proc Natl Acad Sci U S A 2005; 102: 15545–15550.

70 Reich M, Liefeld T, Gould J, Lerner J, Tamayo P, Mesirov JP. GenePattern 2.0 [2]. Nat. Genet. 2006; 38: 500–501.

