## Supplemental Figures and Tables for "Adrenal cortex size, homeostasis and tumorigenesis is regulated by gonadal hormones via androgen receptor/β-catenin signalling crosstalk"

**This PDF file includes:**

Figs. S1 to S8

Table S1, S2

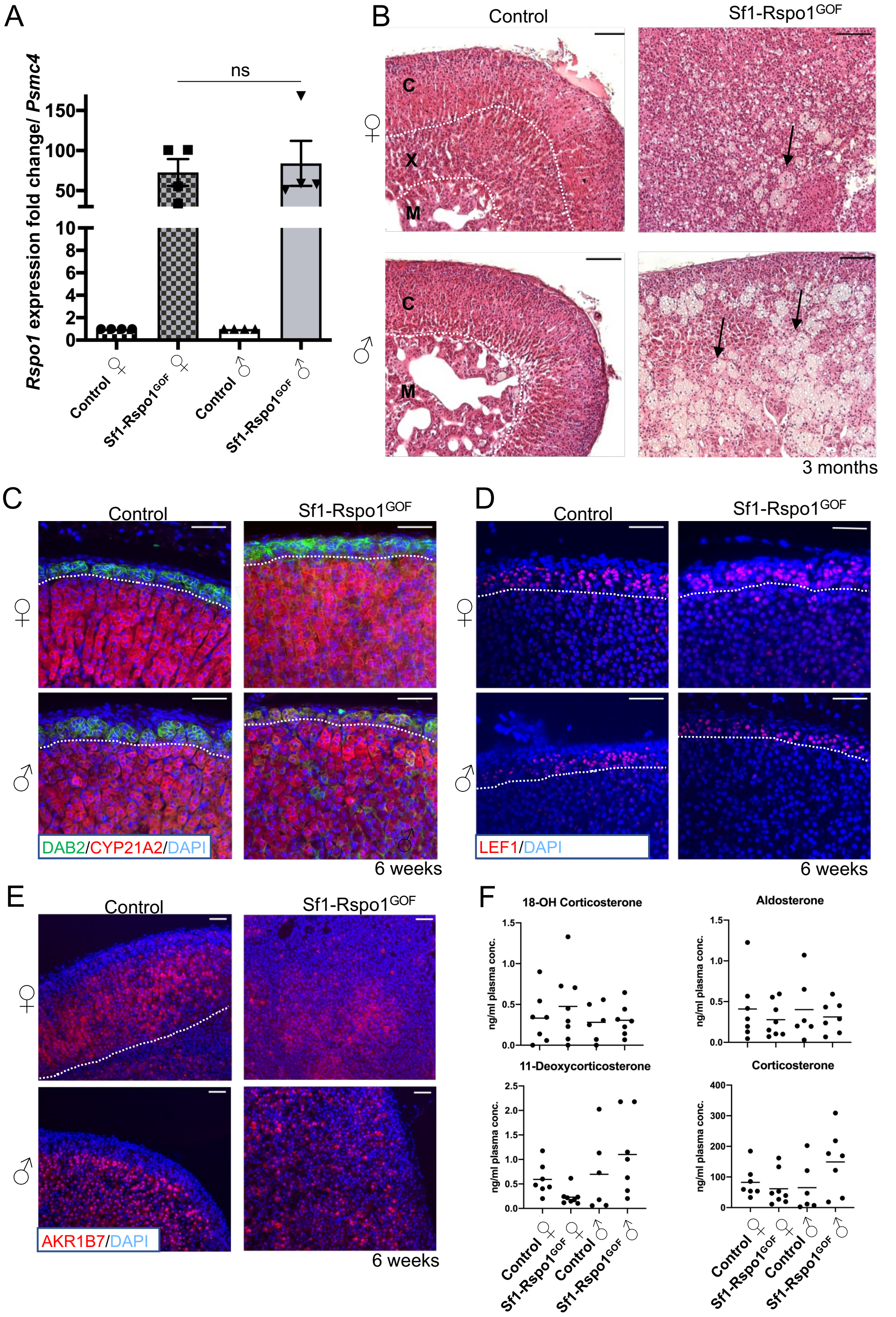
A
B
C
D
E
F
ns
Rspo1 expression fold change/ Psmc4
Control ♀
Sf1-Rspo1GOF ♀
Control ♂
Sf1-Rspo1GOF ♂
Control
Sf1-Rspo1GOF
C
X
M
3 months
DAB2/CYP21A2/DAPI
6 weeks
LEF1/DAPI
AKR1B7/DAPI
18-OH Corticosterone
Aldosterone
11-Deoxycorticosterone
Corticosterone
ng/ml plasma conc.

**Figure S1. Characterisation of RSPO1 overexpression and its effects on adrenocortical cells.** A) RT-qPCR analysis of *Rspo1* expression relative to *Psmc4* in adrenals from 6 week-old mice, shown as mean fold change in *Sf1-Rspo1*$^{GOF}$ adrenals compared to sex-matched controls (n=4). Statistical analysis was performed using unpaired two-tailed t-test (P=0.7442). Error bars represent the standard error of the mean (SEM). B) H&E staining of adrenal sections from 3 month-old mice. Black arrows point to the vacuolated cells forming degenerative lesions. C: cortex, X: x-zone, M: medulla. Scale bar: 100 µm. C) Immunofluorescence staining for DAB2 (marker of the zG) and CYP21A2 (marker of adrenal steroidogenic cells), using adrenal sections from 6 week-old mice. Scale bar: 50 µm. D) Immunofluorescence staining for LEF1 (marker of the zG) on adrenal sections from 6 week-old mice. Scale bar: 50 µm. E) Immunofluorescence staining for the aldo-keto reductase AKR1B7 using adrenal sections from 6 week-old mice. Scale bar: 50 µm. F) Quantification of adrenal steroids in the plasma of control and *Sf1-Rspo1*$^{GOF}$ mice by liquid chromatography paired with tandem mass spectrometry (LC-MS/MS).

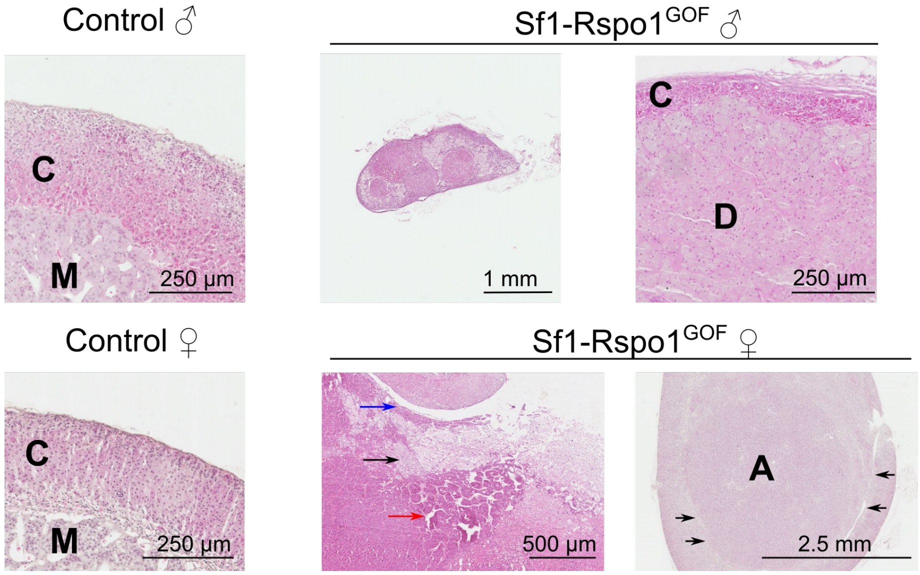


**Figure S2. Ectopic RSPO1 expression leads to the development of nodules and neoplasia in aging animals.** On the top panel, representative H&E staining on 12 month-old male control and *Sf1-Rspo1*$^{GOF}$ adrenals shows the extensive degeneration and cortical thinning paired with incidences of nodular hyperplasia in the *Sf1-Rspo1*$^{GOF}$ adrenals. On the bottom panel, representative H&E images of 12 month-old female control and *Sf1-Rspo1*$^{GOF}$ adrenals are shown. Both female *Sf1-Rspo1*$^{GOF}$ adrenals have tumours: in the middle, the adrenal is completely effaced by an adrenocortical carcinoma. Capsular invasion (blue arrow) and areas of extended necrosis (black arrow) and nest-like cell organisation (red arrow) can be seen. On the left, the adrenal harbours a well-circumcised adrenocortical adenoma. C: cortex, M: medulla, D: degeneration, A: adenoma.

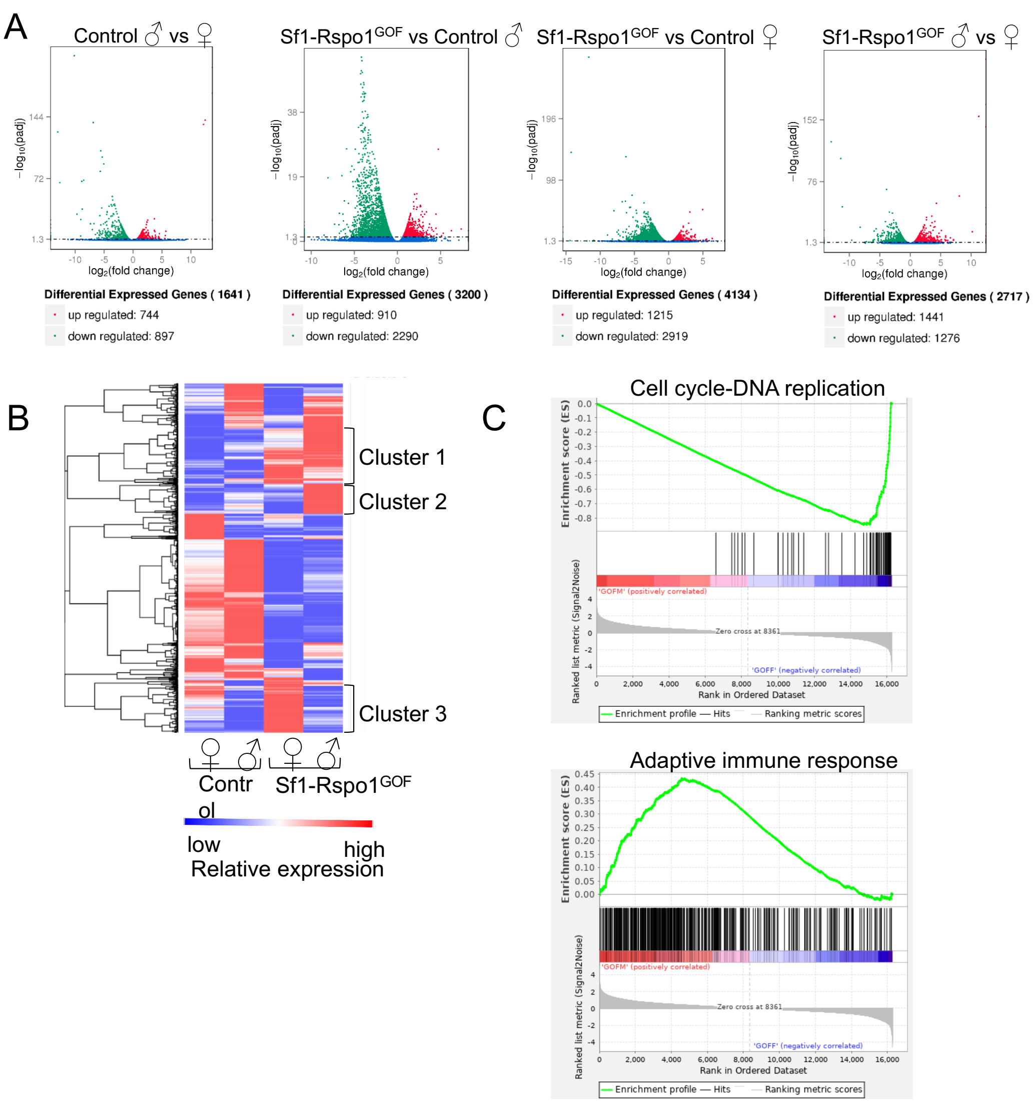


**Figure S3. Analysis of differential gene expression between groups.** A) Volcano plots differentially expressed genes between pairs of experimental groups. B) Hierarchical clustering and heatmap representation of differential gene expression among control and *Sf1-Rspo1*$^{GOF}$, male and female adrenals. Relative expression differences are shown in colour code. C) Enrichment plots for gene ontology (GO) terms produced by GSEA analysis comparing male ('GOFM') to female ('GOFF') *Sf1-Rspo1*$^{GOF}$ adrenals.

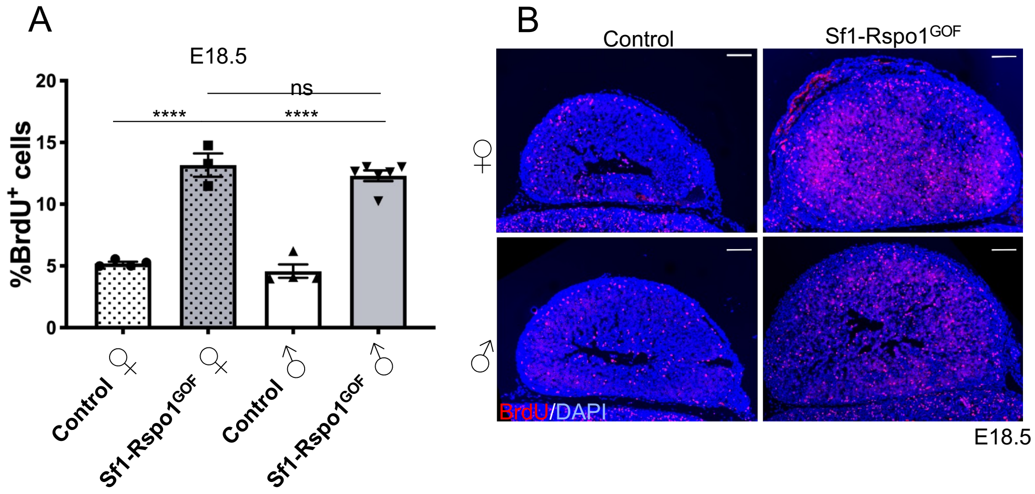


**Figure S4. Male and female Rspo1$^{GOF}$ embryos display equal rates of adrenocortical cell proliferation.** A) BrdU proliferation analysis shown as mean percentage of proliferating cells over total number of cells in the adrenal cortex of E18.5 mouse embryos (error bars represent SEM) (n=3). Statistical analysis was conducted using ordinary one-way ANOVA followed by Tukey's multiple comparison's test. Adjusted p-values: Control F vs *Sf1-Rspo1$^{GOF}$* F: P<0.0001, Control M vs *Sf1-Rspo1$^{GOF}$* M: P<0.0001, *Sf1-Rspo1$^{GOF}$* F vs *Sf1-Rspo1$^{GOF}$* M: P=0.6680. B) Representative images of BrdU immunostaining in E18.5 adrenals. Scale bars: 100 µm.

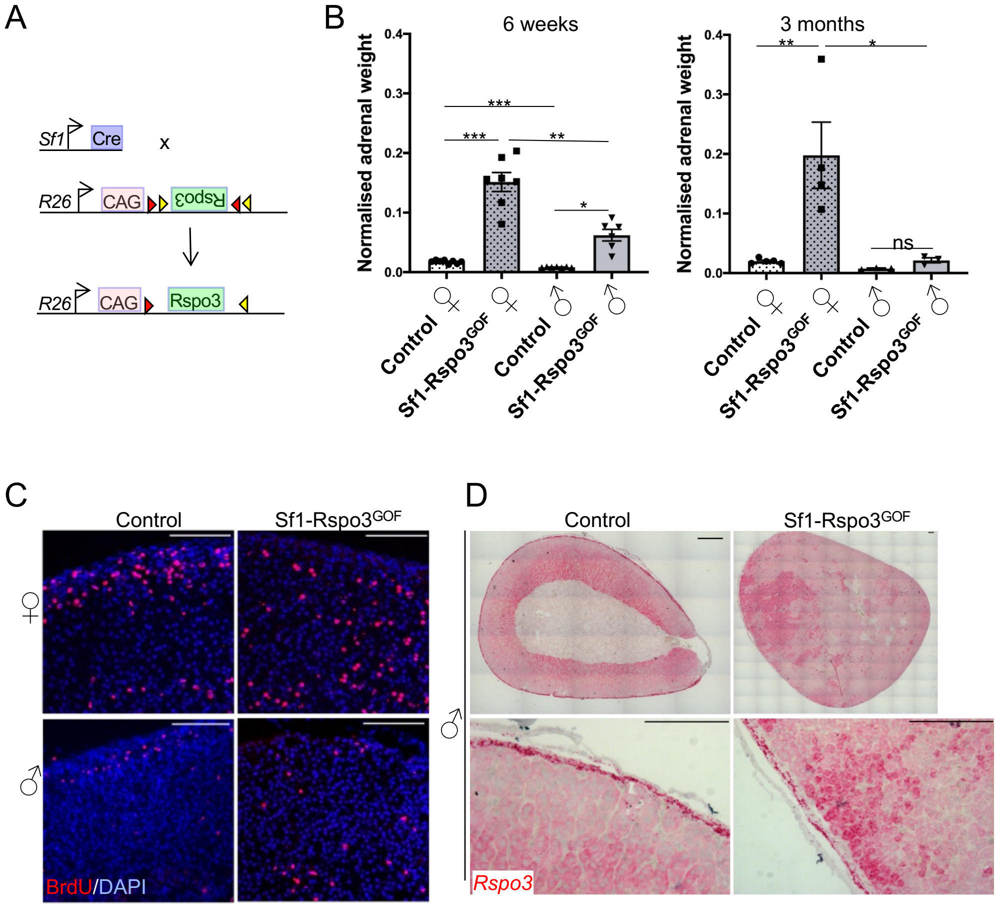


**Figure S5. Ectopic RSPO3 expression results in female-specific hyperplasia.** A) Schematic representation of the genetic strategy to overexpress RSPO3 in adrenocortical cells using Sf1-Cre. B) Mean adrenal weight normalised to total body weight from 6 week and 3 month-old male and female control or *Sf1-Rspo3*$^{GOF}$ adrenals. Error bars represent SEM. Statistical analysis for 6 weeks was conducted using Welch's one-way ANOVA followed by the Dunnett's T3 multiple comparisons test. Adjusted P-values: Control F vs *Sf1-Rspo3*$^{GOF}$ F: P= 0.0008, Control F vs Control M: P=0.0001, *Sf1-Rspo3*$^{GOF}$ F vs *Sf1-Rspo3*$^{GOF}$ M: P=0.0043, Control M vs *Sf1-Rspo3*$^{GOF}$ M: P=0.0130. Statistical analysis for 3 months was conducted with one-way standard ANOVA followed by Tukey's post-hoc test. Adjusted P-values: Control F vs *Sf1-Rspo3*$^{GOF}$ F: P=0.0039, *Sf1-Rspo3*$^{GOF}$ F vs *Sf1-Rspo3*$^{GOF}$ M: P=0.0102, Control M vs *Sf1-Rspo3*$^{GOF}$ M: P=0.9910. C) BrdU analysis to label proliferative cells in control and *Sf1-Rspo3*$^{GOF}$ adrenal at 6 weeks of age. D) In situ hybridisation for *Rspo3* using the RNA Scope method (*Rspo3* mRNA shown as red dots). Representative images represent control and *Sf1-Rspo3*$^{GOF}$ adrenals from male 6 week-old animals. Scale bars: 50 µm.

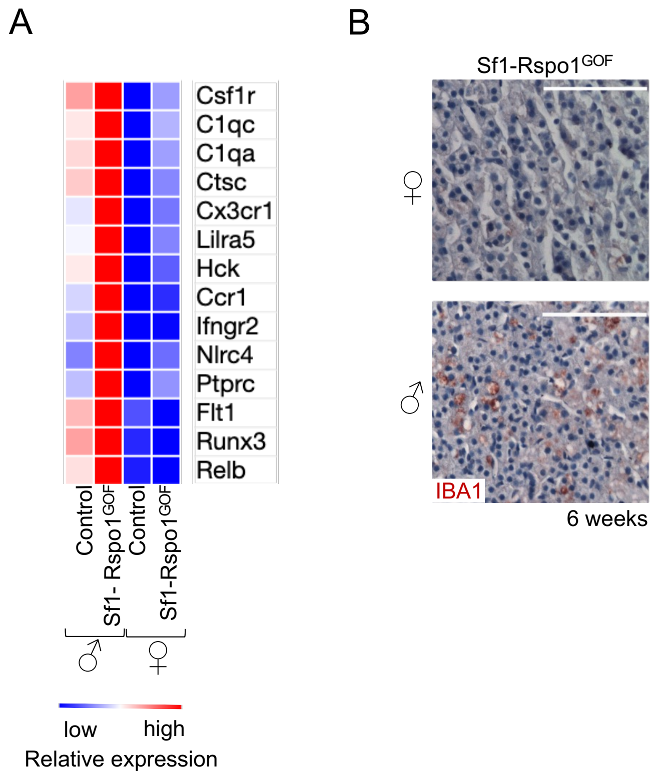


**Figure S6. Degenerative lesion formation by macrophages in the male *Sf1-Rspo1$^{GOF}$* adrenal cortex.** A) Heatmap representation of relative expression differences (shown in colour code) for genes expressed in macrophages/monocytes or pan-immune cells. B) Immunohistochemical staining for IBA1, a macrophage marker enriched in StAR deficient adrenals. Scale bar: 50 µm.

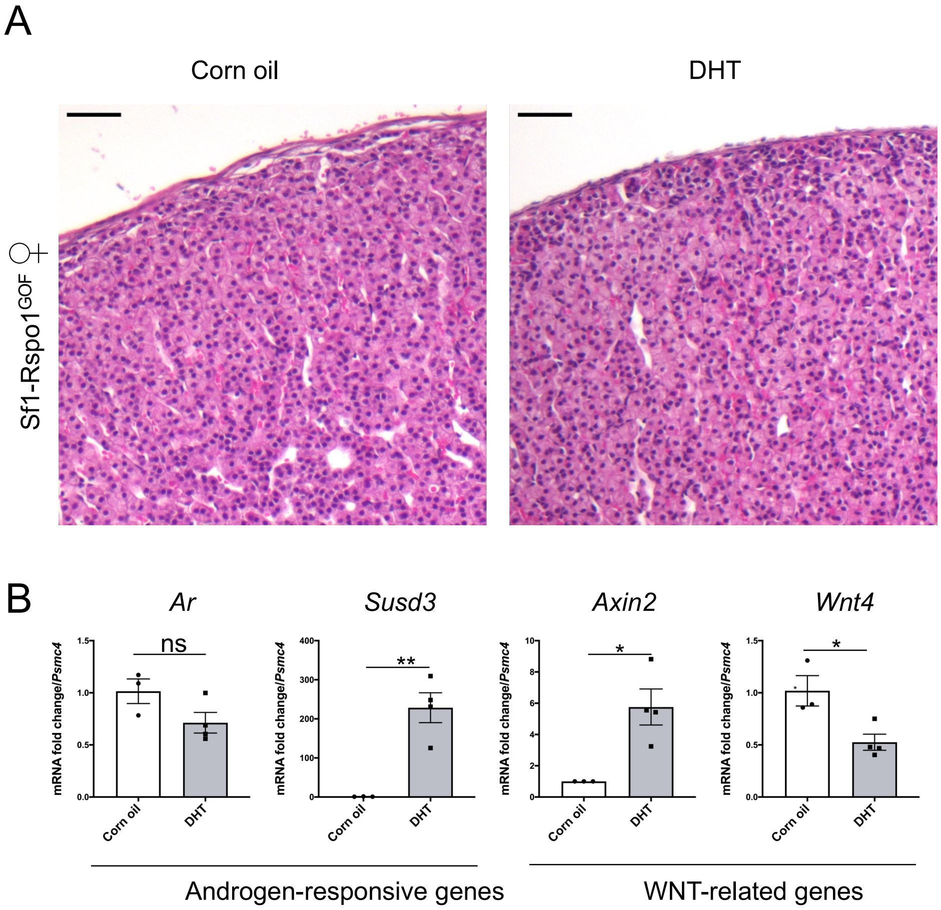


**Figure S7. Effect of DHT treatment on histology and expression in *Sf1-Rspo1$^{GOF}$* adrenals.** A) H&E staining on adrenal sections from 6 week-old female mice treated with corn oil or DHT during puberty. Scale bar: 50 µm. B) RT-qPCR analysis of gene expression for WNT signalling target genes (*Wnt4*, *Axin2*) and genes possibly related to androgen receptor signalling (*Ar*, *Susd3*). Graphs represent mean fold expression change comparing corn oil to DHT treated adrenals (normalised to *Psmc4* expression). Error bars represent SEM. Statistical analysis was conducted with unpaired t-test. P values: 0.1056 (*Ar*), 0.0173 (*Axin2*), 0.0228 (*Wnt4*), 0.0040 (*Susd3*). DHT: dihydrotestosterone.

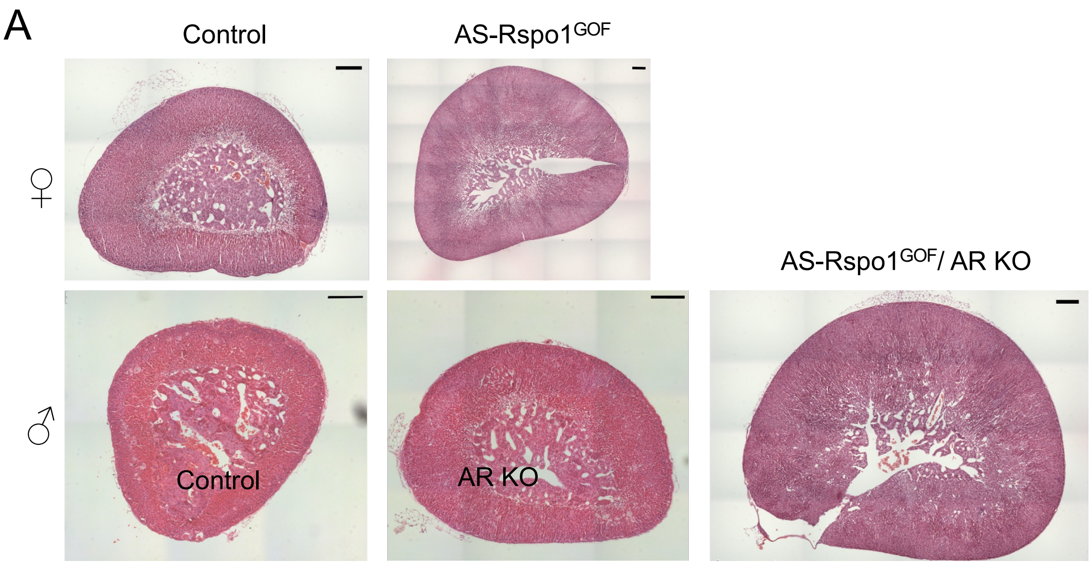


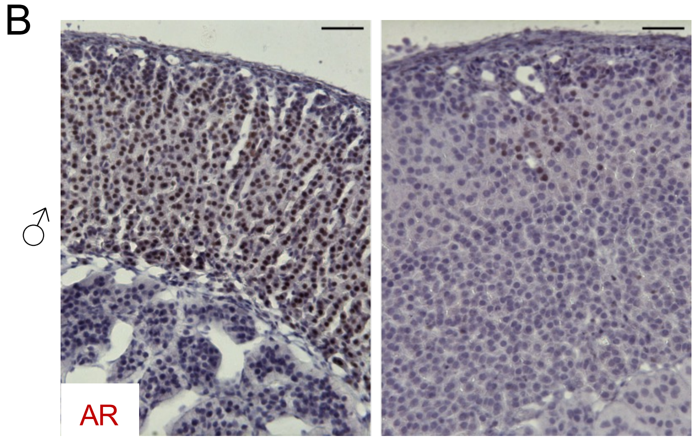


\

**Figure S8: Effect of AR knock-out in adrenocortical histology and AR expression.** A) Hematoxylin and eosin staining of adrenal sections at 20 weeks of age. Scale bar: 200 µm. B) Representative images of immunohistochemistry to detect AR in control and *Ar KO* male adrenals at 20 weeks of age. White arrows point to a few nuclei that display residual expression of AR in the *Ar KO* adrenals. Scale bar: 50 µm.

**Supplementary Table S1: List of primers used in this study**

| Gene name | | | | Purpose |
|---|---|---|---|---|
| *Psmc4* | TTCTTGGAAGCTGTGGATCA | TCAGGATGCGCACATAATAGTT | | qPCR |
| *Rspo1* | CAGTGACTATGCGGCTTGG | AGAGCTCACAGCCCTTGG | | qPCR |
| *Susd3* | GACTGTGCTCATATTCCACTGC | TAAAGCCGAAGGTCTCATGC | | qPCR |
| *Ar* | GTGCCCACCCAAAGGACT | CCTTTTCCCAATGCCTCAG | | qPCR |
| *Pcna* | CTAGCCATGGGCGTGAAC | GAATACTAGTGCTAAGGTGTCTGCAT | | qPCR |
| *Pola1* | TGCACTGTGGATAAATTCTCAAG | CAGTGAGGTTTGACCCATCC | | qPCR |
| *Cdc6* | TCCGTGTGTGGACGTAAAAC | GGAGTGTTGCACAGGTTGTC | | qPCR |
| *E2f8* | CCAGAAATCAGCCCAAACA | GCAGACTGCTCAGCCTCTAAG | | qPCR |
| *Rspo1* transgene | TGTTCATGTCGGGGTTGCGG | CGACCTGCAGCCCAAGCTAG | | Genotyping |
| *Rosa26* | AGGGAGCTGCAGTGGAGTAG | AGCCTGCCCAGAAGACTCCC | | Genotyping |
| Sf1-Cre | CCCACCGTCAGTACGTGAGATATC | CGCGGTCTGGCAGTAAAAACTAT | | Genotyping |
| *Rspo3* transgene | CGCGATTAAATCGATCCCG | CCTATCTGCTTCATGCCAATCC | | Genotyping |
| *Wt1-Sox9* transgene | CATCCGAGCCGCACCTCATG | GCTGGAGCCGTTGACGCG | | Genotyping |
| *Cyp11b2-Cre* | GAGCTGGGGCCCATTTTCAGG | GCTCCAGGTGCATCCGACGG | AACTTGCACCATGCCGCCCA | Genotyping |
| | | AACTTGCACCATGCCGCCCA | | |
| *Sry* | TTGTCTAGAGAGCATGGAGGGCCATGTCAA | CCACTCCTCTGTGACACTTTAGCCCTCCGA | | Genotyping |
| *Ar* flox | AGCCTGTATACTCAGTTGGGG | AATGCATCACATTAAGTTGATACC | | Genotyping |

### Supplementary Table S2: List of primary antibodies used in this study

| Primary antibodies | Source | Reference | Dilution |
|---|---|---|---|
| Anti-Tyrosine hydroxylase (TH) | Sigma-Aldrich | AB152 | 1:200 |
| Anti-CYP21A2 | Sigma-Aldrich | HPA048979 | 1:200 |
| Anti-AKR1B7 (M13) | Santa Cruz | sc-27763 | 1:200 |
| Anti-3BHSD (P18) | Santa Cruz | sc-30820 | 1 :200 |
| Anti-AR | Abcam | ab108341 | 1 :200 |
| Anti-IBA1 | Proteintech | 10904-1-AP | 1 :200 |
| Anti-CD68 | Proteintech | 28058-1-AP | 1:2000 |
| Anti-DAB2 (E-11) | Santa Cruz | sc136964 | 1:50 |
| Anti-BrdU (3D4) | BD Pharmingen | 555627 | 1:100 |
